# ONECUT2 Activates Diverse Resistance Drivers of Androgen Receptor-Independent Heterogeneity in Prostate Cancer

**DOI:** 10.1101/2023.09.28.560025

**Authors:** Chen Qian, Qian Yang, Mirja Rotinen, Rongrong Huang, Hyoyoung Kim, Brad Gallent, Yiwu Yan, Radu M. Cadaneanu, Baohui Zhang, Salma Kaochar, Stephen J. Freedland, Edwin M. Posadas, Leigh Ellis, Dolores Di Vizio, Colm Morrissey, Peter S. Nelson, Lauren Brady, Ramachandran Murali, Moray J. Campbell, Wei Yang, Beatrice S. Knudsen, Elahe A. Mostaghel, Huihui Ye, Isla P. Garraway, Sungyong You, Michael R. Freeman

## Abstract

**Significance Statement:** ONECUT2 (OC2) is a master transcription factor that alters lineage identity by activating gene networks associated with both neuroendocrine prostate cancer and prostate adenocarcinoma. A small molecule inhibitor of OC2 represses the lineage plasticity program activated by enzalutamide, suggesting OC2 inhibition as a novel therapeutic strategy to prevent emergence of treatment-resistant variants.

Graphic Abstract

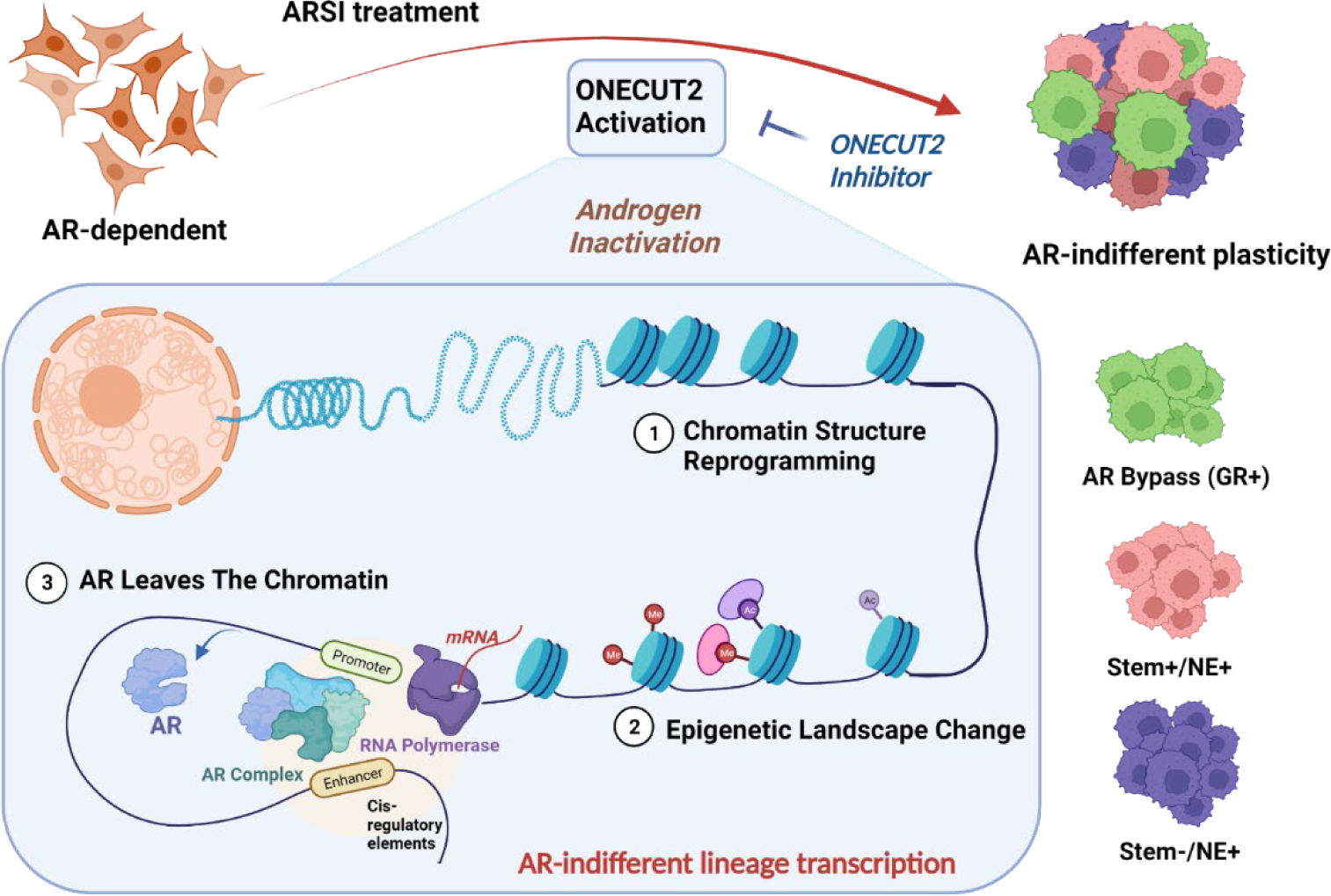

Androgen receptor-(AR-) indifference is a mechanism of resistance to hormonal therapy in prostate cancer (PC). Here we demonstrate that the HOX/CUT transcription factor ONECUT2 (OC2) activates resistance through multiple drivers associated with adenocarcinoma, stem-like and neuroendocrine (NE) variants. Direct OC2 targets include the glucocorticoid receptor and the NE splicing factor *SRRM4*, among others. OC2 regulates gene expression by promoter binding, enhancement of chromatin accessibility, and formation of novel super-enhancers. OC2 also activates glucuronidation genes that irreversibly disable androgen, thereby evoking phenotypic heterogeneity indirectly by hormone depletion. Pharmacologic inhibition of OC2 suppresses lineage plasticity reprogramming induced by the AR signaling inhibitor enzalutamide. These results demonstrate that OC2 activation promotes a range of drug resistance mechanisms associated with treatment-emergent lineage variation in PC. Our findings support enhanced efforts to therapeutically target this protein as a means of suppressing treatment-resistant disease.

## INTRODUCTION

Prostate cancer (PC) is driven by the AR, a hormone-dependent nuclear receptor. AR-driven prostate adenocarcinoma can evolve to contain cell types with diminished luminal features, indicating lineage identity has been altered. This “lineage plasticity” is thought to play a key role in tumor heterogeneity and development of lethal disease. Treatment-resistant phenotypes documented in PC include epithelial-mesenchymal transition (EMT) [1], neuroendocrine (NE) differentiation [2], and activation of the glucocorticoid receptor (GR; *NR3C1*) [3]. While EMT and NE transcriptional programs operate outside the AR axis and give rise to distinct histologic features, the GR assumes control of certain AR-regulated genes, resulting in preservation of the luminal phenotype of adenocarcinoma.

The HOX/CUT protein ONECUT2 (OC2) is a master transcription factor that is active in roughly 60% of mCRPC [4, 5]. OC2 promotes NEPC features, suggesting it plays a role as a driver of lineage plasticity and the emergence of drug resistance following ARSI therapy. Notably, OC2 can be directly targeted with a small molecule inhibitor (SMI) that suppresses established PC metastases in mice [4]. Despite these insights, OC2 activity in disease progression is not well defined. In this study, we describe a novel role for OC2 as a broadly acting lineage plasticity driver that operates across several distinct molecular pathways to promote lineage variation and drug resistance.

## RESULTS

### OC2 is expressed in multiple CRPC lineages

To characterize OC2 expression in CRPC metastases (mCRPC), we performed in-situ hybridization (ISH) of 152 tumor cores from 53 tumors described in a recent study [6]. RNA-seq data published with this series was used to assign AR activity (AR+) status and NE differentiation (NE+) [7] [8], consistent with the published classifications [6]. OC2 mRNA expression was detected in most specimens (52 out of 53) in at least one core (Figure 1A). Based on the published digital spatial profiling (DSP) classification [6], tumors with high OC2 expression exhibited AR-/NE+, double positive (AR+/NE+), double negative (AR-/NE-), and adenocarcinoma (AR+/NE-) phenotypes. All NEPC tumors exhibited relatively high OC2 expression, consistent with the described role of OC2 as an NE driver [4, 5]. However, >80% of AR+/NE-tumors were OC2-positive, and high OC2 expression was also seen in AR+/NE+ and AR-/NE-specimens (Figure S1A). Notably, 46 AR+/NE-cores displayed high OC2 staining via ISH, which represents 40% of the adenocarcinoma specimens. High ISH staining correlated significantly with high immunohistochemical (IHC) score (Figure S1B). Representative images IHC staining in CRPC with distinct phenotypes are shown in Figures 1B and S1C. These findings indicate that OC2 expression is not restricted to NEPC but can occur widely within a range of CRPC phenotypes, including in tumors with AR pathway activation.

**Fig 1.**
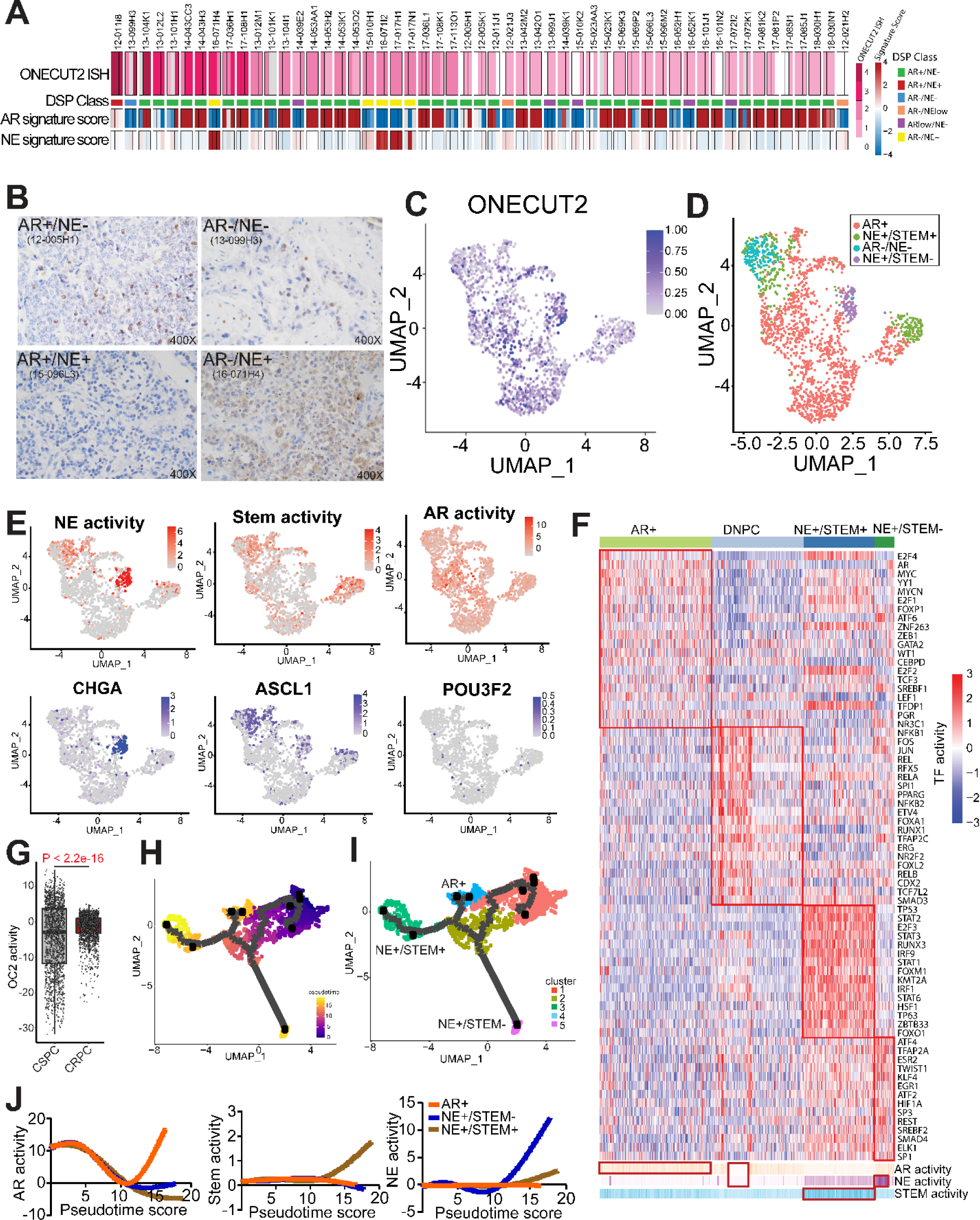
OC2 is expressed in multiple CRPC lineages. (A) RNA scope in-situ hybridization of OC2 on UW-TMA95 series ([6]). NE and AR signature scores were calculated using RNA-seq data and DSP class from the matched samples collected from GSE147250. (**Refer to methods)** (B) Representative IHC staining of OC2 in CRPC specimens with distinct features. (C) UMAP plot illustrating OC2-expressing epithelial cells from 6 CRPC patients scRNA-seq datasets selected for further analysis. (D) UMAP plot illustrating OC2-expressing cells annotated to distinct lineages in CRPC. (E) UMAP plots illustrating OC2-expressing cells colored by three lineage signature activities (AR, NE, Stem) and expression levels of representative markers (*CHGA, ASCL1 and POU3F2*). (F) Unsupervised clustering of OC2 expressing epithelial cells from 6 CRPC scRNA-seq datasets using transcription factor (TF) activity from the high-confidence DoRothEA database. (**Refer to methods**) (G) CRPC patient samples show higher OC2 activity (**Refer to methods**) compared to CSPC patient samples. (H) UMAP plot showing the pseudotime trajectory of OC2-expressing cells with progression from CSPC (M0 status) cells to CRPC. (I) UMAP plot showing development of distinct lineages along the pseudotime trajectory. (J) Trajectory of development of 3 lineages.

To further investigate these diverse lineages and OC2 expression, we employed single-cell RNA sequencing (scRNA-seq) data from three independent studies comprising 14 castration-sensitive PC (CSPC) and six patients with CRPC [9–11]. We performed data-specific bias correction, clustering, and access on cell data with epithelial marker gene expression (Figure S1E) (see details in Methods). OC2-expressing epithelial cells were then selected from CRPC data in the combined scRNA-seq data (Figure S1E, Figure 1C) for further analysis. Markers associated with distinct phenotypes in OC2-expressing epithelial cells were displayed using Uniform Manifold Approximation and Projection (UMAP) plots (Figure 1D) and lineage signatures were mapped to the OC2-expressing cells (Figure 1E). Notably, AR activity was evident in most cells. The neural differentiation indicator chromogranin A (CHGA) overlapped with the NE+/Stem-phenotype, while the NE driver ASCL1 overlapped with both NE+/Stem- and NE+/Stem+ phenotypes, consistent with descriptions of the role of ASCL1 in CRPC [12]. Expression of the NE master regulator BRN2 [13] was too low to be detected at the single-cell level (Figure 1E). These results, where NE+ cells cluster into two different groups, are consistent with the results of the Brady et al. study [6] (Figure S1D).

To address this heterogeneity, we computed 118 TF activity profiles based on mRNA expression of known targets, encompassing all high confidence targets [14] in the OC2-expressing epithelial cells.Four TF modules were identified through hierarchical clustering [15] (Figure S1F), which correspond to distinct lineages based on standard markers representing AR, Stem and NE features. Four subtypes expressing OC2 were subsequently defined as displaying AR+, AR-/NE-, NE+/Stem+, and NE+/Stem-phenotypes (Figure 1F). Consistent with recent reports, the phenotype exhibiting Stem+ displays elevated interferon signaling (Figure S1G), with high activity of IRF1/9 and STAT1/2/3/6 [16]. Taken together, these results demonstrate that OC2-positivity coincides with distinct molecular phenotypes aligned to modules defined by divergent TF activity.

OC2 activity computed by the gene signature [4] was significantly higher in CRPC compared with CSPC (Figure 1G). In CSPC, M1 metastatic disease in CSPC patients displayed higher OC2 activity in comparison to M0 patients (no detectable metastases). Similar results were seen in bulk RNA-seq data of primary tumors in M0 and M1 patients [17] (Figure S1H). These results suggest that OC2 activity correlates with more advanced disease. To track the progression trajectory of OC2-expressing epithelial cells, we first separated CSPC and CRPC cells in the combined scRNA-seq data. Pseudotime analysis from CSPC (M0 status) to CRPC revealed three branches developing during progression to CRPC (Figure 1H-I). Established signatures were applied to assess pseudotime-dependent changes of AR+, NE+/Stem- and NE+/Stem+ lineages (Figure 1J). These suggest increases of NE+ and Stem+ activity at two terminal points (NE+/Stem+ and NE+/Stem-clusters). Consistent with this, OC2 activity is also high in the same terminal points (Figure S1I-J). Prostate stem cell antigen (PSCA) is highly expressed at the starting point (CSPC), while ASCL1 expression is enriched in NE+/Stem+ and NE+/Stem-clusters (Figure S1J). PSCA is reported to be a marker for Stem-like Luminal L2 cells, described as a source of emergent lineage plasticity [18]. These findings show OC2-expressing cells progressing along a trajectory from CSPC appear to adopt multiple routes to bypass AR dependence.

### OC2 alters chromatin accessibility

To recapitulate the progression from CSPC to CRPC, OC2 was enforced in AR-dependent LNCaP cells. To uncover the mechanism underlying OC2-induction of heterogeneous lineages, we investigated the OC2 interactome using Immunoprecipitation-Mass Spectrometry (IP-MS). The most abundant peptides derived from the OC2 interaction trap represented proteins involved in chromatin remodeling processes (Figure 2A; Figure S2A). To determine whether OC2 expression contributes to global chromatin remodeling, we performed Hi-C chromosome capture [19, 20] and integrated these data with H3K27 acetylation (H3K27Ac) profiles. OC2 OE increased enhancer-promoter interactions genome-wide, suggesting the newly formed looping structures contributed to OC2-directed gene expression changes (Figure 2B). The endogenous OC2 gene locus also showed specific enhancers looped to the OC2 promoter in OC2 OE LNCaP cells (Figure S2B). This led us to assess chromatin accessibility across the genome in OC2 OE cells using transposase-accessible chromatin with high-throughput sequencing (ATAC-seq). Genome-wide chromatin accessibility was increased in OC2 OE cells (Figure S2C-D), with 40,568 unique regions opened under this condition (Figure 2C). TF motif analysis of OC2-induced hyper-accessible regions revealed enrichment of FOXA1, HOXB13, GRE (glucocorticoid receptor element), AR-half-site and NANOG motifs (Figure 2D). To decipher the possible role of OC2 in chromatin remodeling, the OC2 cistrome was profiled with CUT&RUN-seq under the same conditions. Known TF binding motifs were ranked by similarity score to distinct OC2 binding sites. OC2 OE cells exhibited a distinct TF binding profile in comparison to control cells. As previously shown [4], in non-perturbed cells OC2 shows co-occupancy with the AR, whereas YY1 was the top co-occupied TF in OC2 OE cells (Figure 2E). Intriguingly, ∼ 75% of the OC2-bound regions consisted of closed chromatin. OC2 OE resulted in an increase in the number of OC2 binding peaks at closed chromatin. The ratio of closed/open peaks with OC2 binding increased from 2.5 (19,770 vs. 8,134) to 3.5 (31,916 vs. 9,216) with enforced OC2 (Figure 2F), suggesting a chromatin remodeling function.

**Fig 2.**
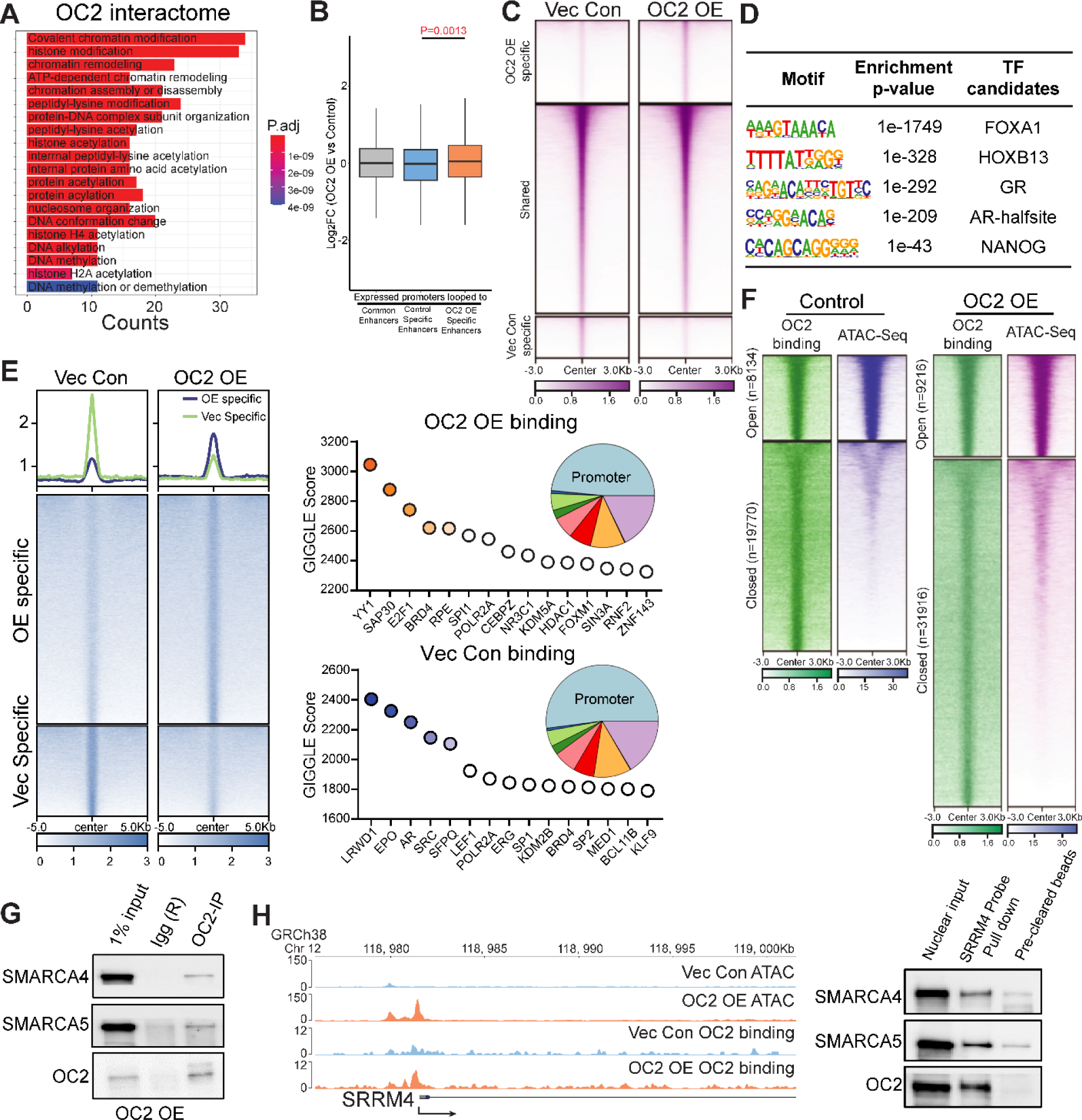
OC2 interacts with CRCs and alters chromatin accessibility. (A) Gene Ontology analysis of the OC2 interactome identified by IP-MS experiments (N = 2). (B) Integrated analysis of HiC-seq, RNA-seq and histone H3K27Ac CUT&RUN-seq showed activated gene promoter regions looped to OC2 OE specific enhancers. (C) Normalized tag densities for ATAC-seq in LNCaP Vec Con and OC2 OE cells. (N = 2) (D) Motif enrichment analysis for hyper-accessible regions in OC2 OE from ATAC-seq data. (E) Normalized tag densities for the OC2 cistrome in LNCaP Vec Con and OC2 OE cells through CUT&RUN-seq (Left). GIGGLE analysis for TF binding similarity in the OC2-specific cistrome in both conditions (Right). (N = 2) (F) Normalized tag densities for integrated analysis of ATAC-seq and OC2 CUT&RUN-seq. 75% of the OC2-bound regions consisted of closed chromatin. (G) IP-WB showed OC2 interacts with SMARCA4 and SMARCA5. (H) DNA affinity precipitation assay (DAPA) showed binding of SMARCA4 and SMARCA5 to the SRRM4 promoter region under OC2-enforced conditions. (**Refer to methods**)

Chromatin remodeling complexes (CRCs) modify chromatin architecture to allow TF access to condensed genomic DNA [21]. Four CRC families exist in humans: Chromodomain Helicase DNA-binding (CDH), INO80, SWItch/Sucrose Non-Fermentable (SWI/SNF) and Imitation switch (ISWI) complexes. OC2 interactome data indicate that OC2 physically associates with multiple proteins in CRCs representing the SWI/SNF and ISWI complexes (Figure S2E). The most abundant CRC peptides identified through IP-MS were those of SMARCA4 (SWI/SNF) and SMARCA5 (ISWI). INO80 and CDH families were either low in abundance or undetectable.

OC2 complexed with SMARCA4 and SMARCA5 (Figure 2G), demonstrating that both proteins interact with endogenous OC2. To explore the interaction of OC2 with CRCs at a physiologically relevant locus, the serine/arginine matrix 4 (SRRM4) promoter was selected. SRRM4 is an NE-associated splicing factor shown to be a lineage plasticity driver in CRPC [22]. The OC2 OE condition resulted in an increase in both SRRM4 mRNA level and chromatin accessibility at the endogenous SRRM4 promoter. OC2 bound to the SRRM4 promoter, consistent with OC2 as a direct regulator of this gene. OC2, SMARCA4 and SMARCA5 bound to a cloned segment of the SRRM4 promoter (Figure 2H). These findings confirm that OC2 interacts with chromatin remodeling proteins at a genomic locus where its enforced expression exerts a robust remodeling effect.

### OC2 activates multiple AR-independent lineage-defining factors

We found multiple lineage-defining TF motifs were newly accessible under OC2 OE conditions (Figure 2D). To identify TF drivers and their transcriptional programs, we performed RNA-seq and ATAC-seq with OC2 OE and control cells. Integrative analysis of global transcriptional and chromatin landscape changes induced by OC2 allowed assessment of differential TF expression, as well as enrichment of TF motifs ion the accessible chromatin regions and downstream target expression levels. We applied TF target gene enrichment (fold enrichment score), where the higher score represents a greater percentage of target gene changes (Figure S3D). This revealed a series of CRPC- and/or NEPC-associated TF genes, including *NR3C1*, *ETV4*, *TWIST1*, *POU3F2*, *TFAP2A*, and *KLF5*, which were highly activated by OC2 OE [3, 13, 23–26].

We determined whether OC2-induced TF activation is associated with epigenetic modifications. OC2 binding ±2kb around transcription start sites (TSS) were evaluated for H3K4me3 and H3K27me3 histone marks, indicating active or repressed epigenomic states, respectively. OC2 OE resulted in a global shift from repressive or bivalent status toward transcriptional activity (Figure 3B, Top). OC2-regulated TFs (marked red in the figure) exhibited an increase in the H3K4me3 activation mark and a decrease in the H3K27me3 repressive mark (Figure 3B, Bottom). In contrast, AR downstream target genes (blue) moved in the opposing direction, indicating AR suppression by OC2 (Figure 3B, Bottom). The cut-off of H3K4me3 signal is based on two-normal distribution (Figure S3E). H3K27me3 and H3K4me3 marks were aligned with ATAC-seq profiles at genomic loci corresponding to AR+, NE+/Stem-, and NE+/Stem+ lineage-defining factors, including those involved in NEPC (POU3F2, SYP), AR bypass (NR3C1, AR), as well as Stem-like features (SOX2, CD44) (Figure 3C). Highly ranked TFs induced by OC2 in LNCaP cells also showed high expression in OC2-high CRPC tumors from the SU2C cohort [27] (Figure 3D). Expression of OC2-regulated TFs correlated with OC2 activity computed by PC OC2 signature [4] in the SU2C CRPC cohort (Figure 3E). We validated that OC2 OE increased the expression of AR-indifferent, lineage-defining factors in another CSPC model, LAPC4 (Figure 3F).

**Fig 3.**
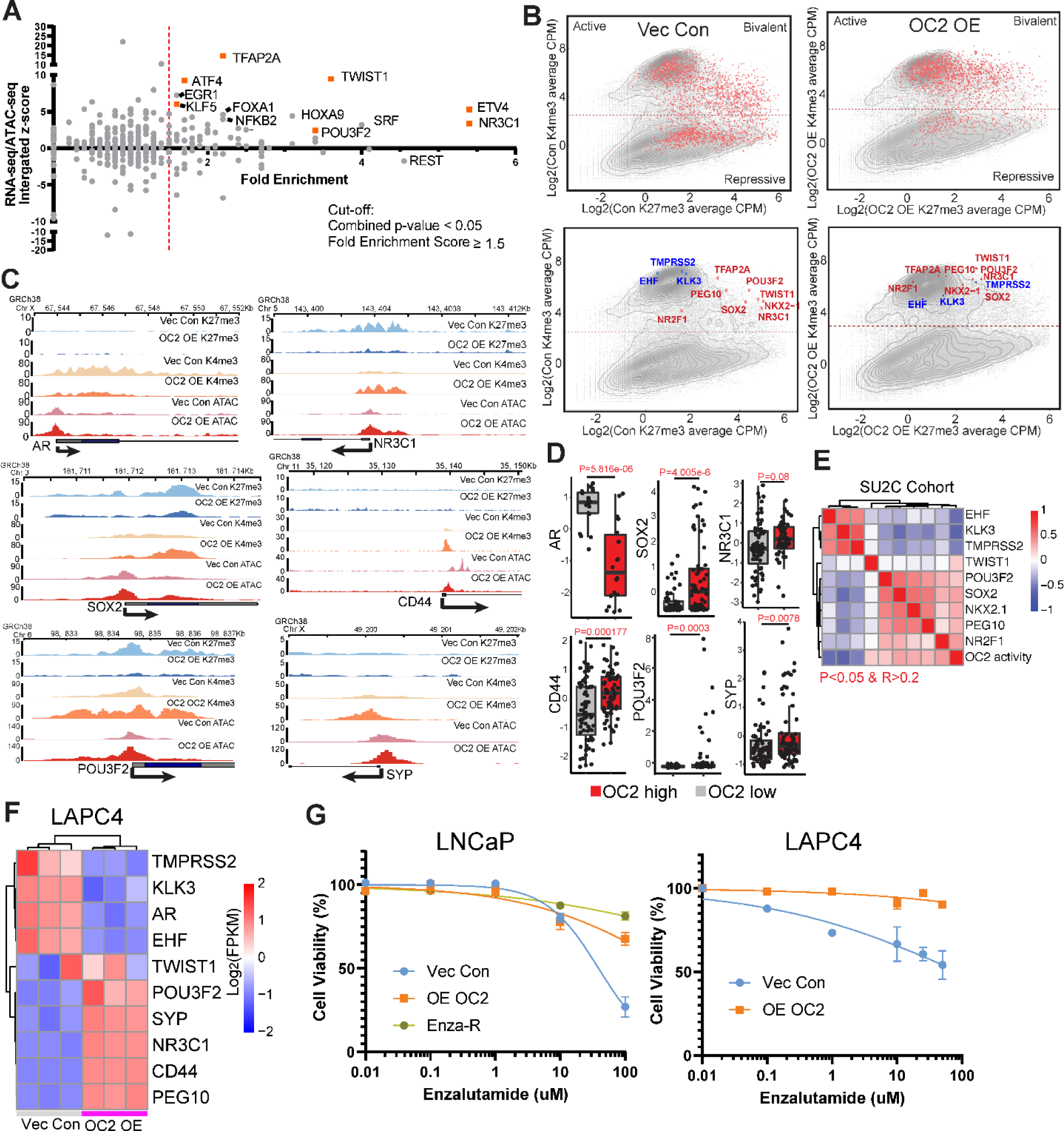
OC2 activates multiple AR-independent lineage-defining factors. (A) Integrated analysis for identifying OC2-driven candidate TFs based on RNA-seq, ATAC-seq and fold enrichment of downstream targets. (**Refer to methods**) (B) Log_2_ (CPM+0.1) signal intensity of H3K4me3 and H3K27me3 within ±2 kb of the transcription start site (TSS) where OC2 binds. Each dot represents a unique transcript start site. The cutoff for H3K4me3 separation is based on two normal distribution of the signal. Selected genes are highlighted in the bottom row. (C) Visualization of H3K27me3, H3K4me3 and ATAC-seq signals on selected genes (AR Bypass: AR/GR, NEPC: POU3F2/SYP, Stem: SOX2/CD44). (D) The expression levels of matched selected genes in the SU2C CRPC cohort based on stratification of OC2 activity. (E) Correlation of OC2 activity with candidate TF expression in the SU2C cohort. (F) OC2-driven TFs were also activated in LAPC4 Vec Con and OC2 OE cells. (N = 3 biologically independent experiments) (G) OC2 overexpression promotes enzalutamide resistance in LNCaP and LAPC4 cells. LogIC50: LNCaP (1.59 (Vec) vs. 2.75 (OC2-OE) vs. 4.46 (Enza-R)); LAPC4 (1.95 (Vec) vs. 5.49 (OC2_OE))

Enforced OC2 OE in LNCaP and LAPC4 models resulted in an enzalutamide-resistant phenotype in both cell lines (Figure 3G). Taken together, these findings indicate that OC2 operates globally at the chromatin level to activate numerous AR-indifferent, lineage-defining factors. Notably, OC2 overexpression alone resulted in the activation of multiple AR-indifferent lineages without ARSI challenge.

### OC2 regulates AR target genes by upregulation of the glucocorticoid receptor

We used the H3K27me3 and H3K4me3 histone marks to investigate the ability of OC2 to regulate the epigenetic status of the promoters of a panel of lineage signature genes. Upon OC2 activation, NE and Stem-like lineage signature scores moved in the direction of epigenetic activation, whereas the AR-governed lineage was repressed (Figure 4A). OC2 OE suppressed prostate specific antigen (PSA/KLK3) mRNA and other AR targets, while OC2 knockdown activated AR signaling (Figure S4A-B). However, residual PSA expression remained even under enforced conditions (Figure 4B). Surprisingly, OC2 silencing in the OE condition, using either siRNA or OC2 inhibitor CSRM-617 [4], suppressed PSA expression instead of restoring it (Figure 4B). A similar result was seen with the AR-target genes TMPRSS2, FKBP5, NKX3.1, and STK39 (Figure S4C). This notable and unexpected finding suggests that OC2 may be capable of activating certain AR-dependent genes.

**Fig 4.**
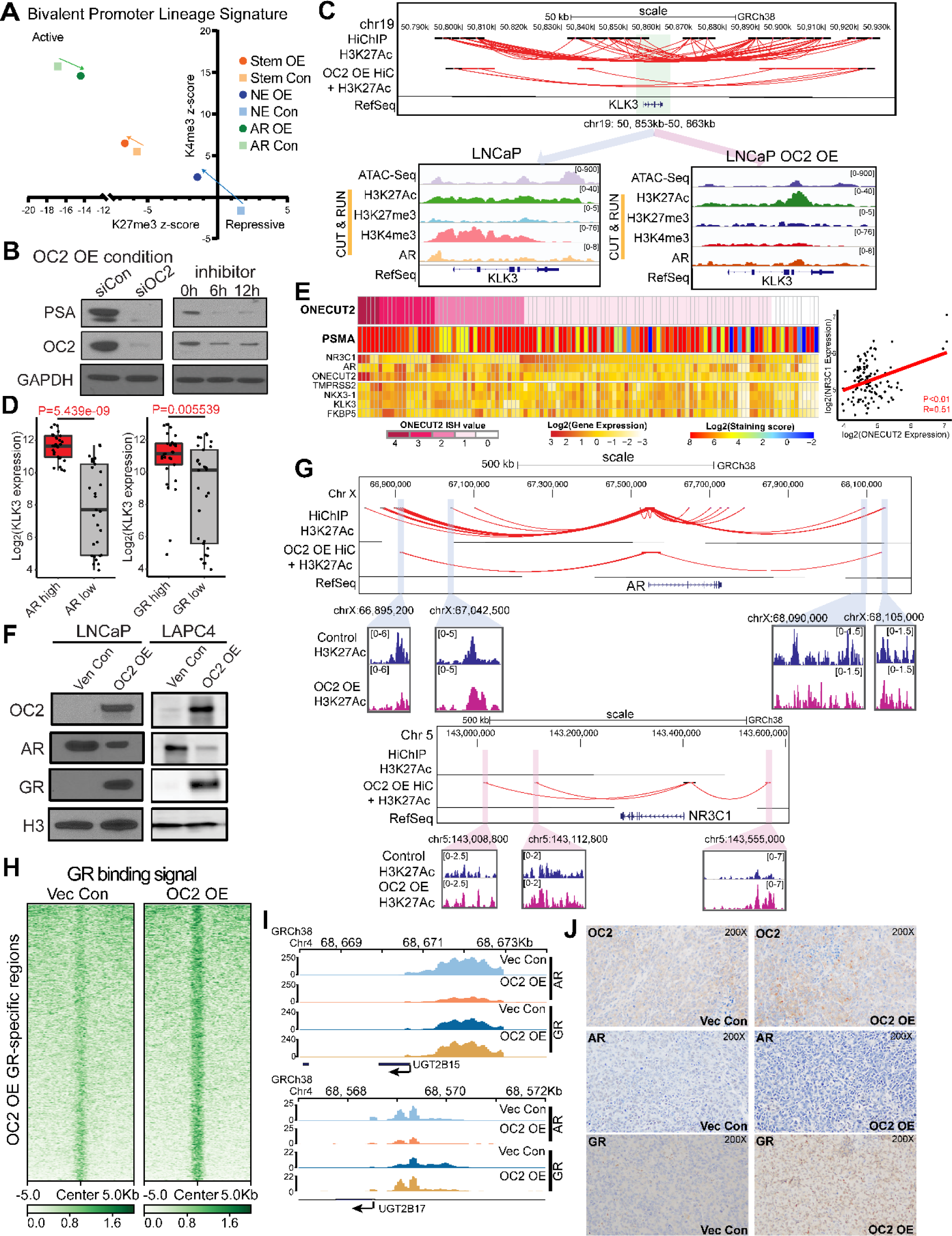
OC2 activates AR-bypass pathways through GR. (A) Epigenetic status (H3K27me3 and H3K4me3 signal) of the promoters of a panel of lineage signature genes (AR/Stem/NE) represented by z-score in LNCaP Vec Con and OC2 OE cells. (B) OC2 controls PSA-expressing lineage in OC2 OE cells. (C) Integrated analysis showed loss of enhancer looping to the *KLK3* promoter (Top). ATAC-seq and CUT&RUN-seq of H3K27Ac, H3K27me3, H3K4me3 and AR binding signal at *KLK3* in LNCaP Vec Con and OC2 OE cells (Bottom). (D) KLK3 expression in AR and GR high vs. low groups in SU2C cohorts. (E) mCRPC specimens [6] with evidence of AR activity, based on AR signature and positive PSMA staining, which show high OC2 expression, also exhibited either high AR or high GR expression (Left). OC2 mRNA level is positively correlated with GR mRNA (Right). (F) OC2-induced GR upregulation with AR suppression in both LNCaP and LAPC4 cells. (G) Integrated HiC with H3K27Ac loops near *AR* and *NR3C1* (GR) promoter loci. (H) Normalized tag densities of CUT&RUN-seq at specific GR binding regions. (I) Loss of AR binding and gain of GR binding at the promoter loci of *UGT2B15* and *UGT2B17* in LNCaP OC2 OE cells. (J) OC2, AR and GR IHC staining in xenograft models created by subcutaneous injection of LNCaP Vec Con and OC2 OE cells.

The suppressive effect of OC2 OE on PSA/KLK3 was seen in the integrated HiC and H3K27Ac data, where substantially fewer chromatin loops were evident in the OC2 OE condition at the *KLK3* promoter and surrounding regions (Figure 4C). This coincided with lower H3K4me3 signal (Figure 4C). Chromatin accessibility and AR binding peaks were also decreased, suggesting that PSA expression is no longer under the control of AR.

One of the top OC2-driven TFs (Figure 3A) is the GR(*NR3C1*), and GREs were identified among the most frequently co-occupied sites with enforced OC2 (Figure 2E). GR can drive AR target gene expression in PC [3]. KLK3 expression was higher in the GR-high subset of the SU2C CRPC cohort contrasted with the GR-low subset (Figure 4D). Similarly, in the Brady et al. cohort [6], AR-positive specimens with AR-driven PSMA staining and high OC2 expression also exhibited either high AR or high GR, with the highest levels of OC2 expression exhibiting high GR expression (Figure 4E). In these specimens, AR and GR expression levels were inversely related, consistent with the ability of AR to suppress GR [28]. High OC2 ISH values coincided with high OC2 mRNA in the Brady et al. cohort (Figure 4E). In both LNCaP and LAPC4 CSPC models, OC2 OE induced GR expression and suppressed AR in the nucleus (Figure 4F). Using an inducible system in LAPC4, GR expression was suppressed when OC2 expression was repressed (Figure S4D), suggesting the GR is directly regulated by OC2. Consistent with this, endogenous OC2 bound to the *NR3C1* promoter (Figure S4E), and the OC2 OE condition significantly increased chromatin accessibility at the promoter (Figure 3C). HiC revealed that OC2 OE resulted in more three-dimensional genomic contacts linked to the *NR3C1* promoter, coinciding with elevated H3K27ac signal in three putative enhancer regions, thus indicating enhancer reprogramming. Chromatin loops to the *AR* promoter were lost (Figure 4G). OC2 OE substantially increased GR binding genome-wide (N= 22,751) as shown by CUT&RUN-seq (Figure 4H). The AR-driven, androgen-inactivating genes *UGT2B15* and *UGT2B17* are upregulated with androgen deprivation therapy or ARSI treatment [29, 30]. AR binding to the *UGT2B15* and *UGT2B17* promoters under OC2 OE was reduced, while GR binding was increased (Figure 4I). In LNCaP xenografts, OC2 OE cells showed reduced AR coinciding with positive GR staining (Figure 4J). Collectively, these results suggest that OC2-driven GR activation restores expression of AR target genes. This phenomenon is associated with ARSI resistance in adenocarcinoma.

### OC2 promotes NE features through super-enhancer reprogramming

OC2-associated genes in CSPC cells were identified in LNCaP OC2 OE cells, with the criterion that they were oppositely regulated under OE and knockout conditions (Fold change > 1.5 and adj.p < 0.05). NE and Stem-like pathways were prominent among the upregulated genes. Repressed pathways include androgen response and P53 hallmarks, suggesting emergence of lineage plasticity (Figure 5A). Enforced OC2 in LNCaP and LAPC4 cells activated a NE differentiation RNA signature [7] seen in aggressive PC (Figure 5B; Figure S5A). Epigenetic activation of ASCL1 is also seen with OC2 OE cells (Figure S5B). Consistent with this, a xenograft model derived from subcutaneous injection of OC2 OE cells exhibited positive SYP staining (Figure 5C). This OC2 activity signature, and a published NEPC signature [31], were applied to 18 patient-derived xenografts [7]. Both signatures comparably identify NE tumors within the cohort, reinforcing the role of OC2 as a promoter of NE differentiation (Figure S5C).

**Fig 5.**
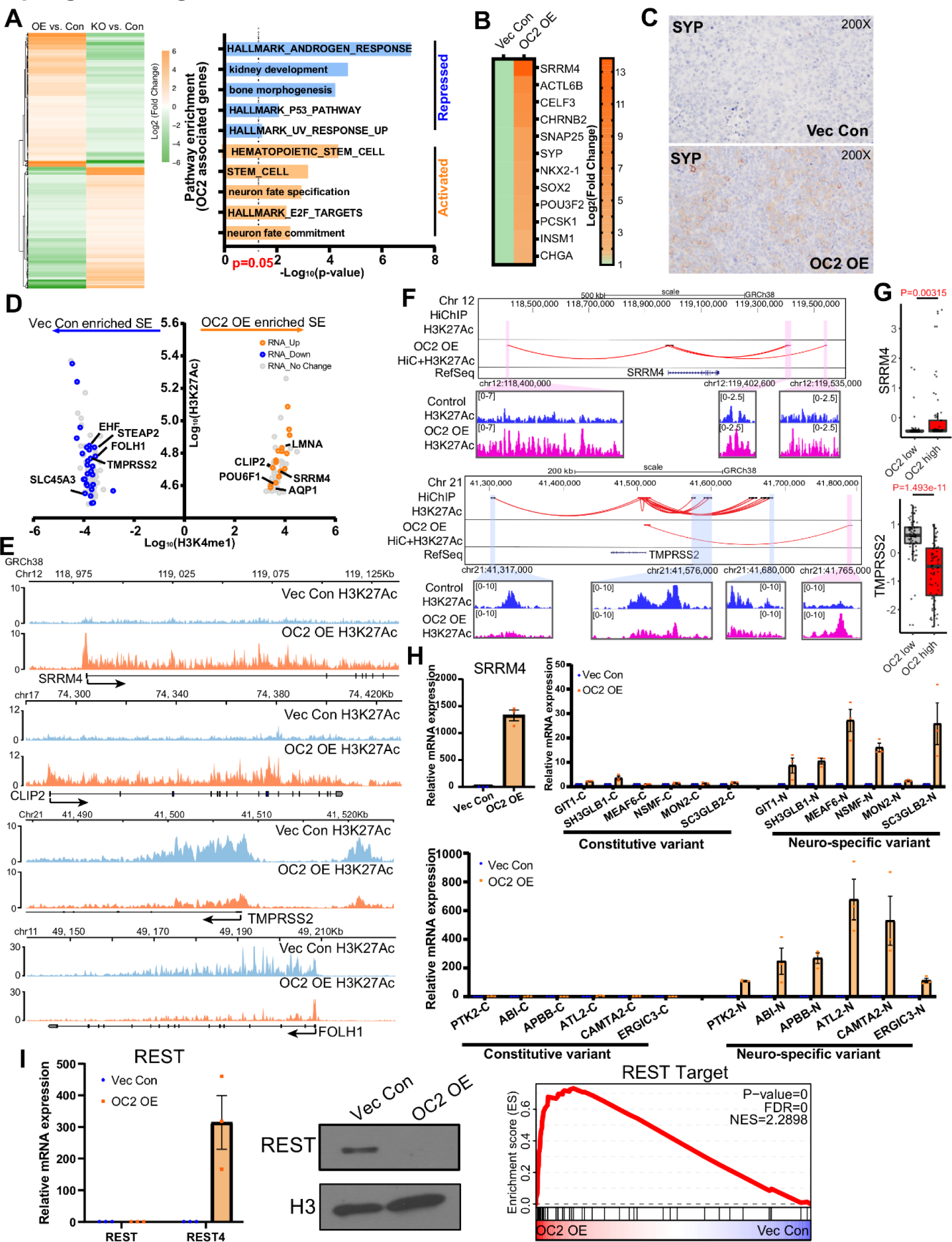
OC2 promotes neuroendocrine features through super-enhancer reprogramming. (A) OC2 signature genes derived from AR-dependent LNCaP cells (Left). And the Geneset enrichment analysis of OC2 signature genes (Right). (B) Enforced OC2 expression upregulates multiple NE signature genes in LNCaP cells. (C) SYP staining in LNCaP Vec Con and OC2 OE cells xenograft models. (D) Genes proximal to super-enhancer regions identified as newly formed in OC2 OE cells vs. control cells with corresponding change in RNA level. (E) Visualization of H3K27Ac signals around *TMPRSS2 / FOLH1* (encoding PSMA) and *SRRM4 / CLIP2* are shown. (F) Integrated HiC and H3K27Ac signal showed enhancers looped to promoter loci of *TMPRSS2* and *SRRM4*. (G) Expression of TMPRSS2 and SRRM4 in SU2C cohorts stratified by OC2 activity. (H) Upregulation of the splicing factor SRRM4 promotes neural-specific splicing variants with OC2 overexpression. (I) Master neuronal suppressor REST was suppressed in the nucleus through SRRM4-mediated spicing variant REST4, which promotes NE differentiation.

Genomic profiling of H3K27Ac was performed in OC2 OE and vector control cells to identify transcriptionally active regulatory elements. The H3K27ac profile was substantially changed genome-wide in response to enforced OC2. Super-enhancers (SEs) are large chromatin domains, densely occupied by H3K27Ac and H3K4me1 enhancer marks, which regulate lineage-specific gene expression [32, 33]. Genes proximal to SE regions identified as newly formed in OC2 OE *vs* control, in which there was a corresponding change in RNA expressed from these genes, are shown (Figure 5D, Figures S5D. and Figures S5E). The most highly upregulated gene with enforced OC2 was *SRRM4* (Figure 5B), a splicing factor and NE-driver shown above to be directly regulated by OC2 (Figure 2H). *SRRM4* is notable as an SE-driven gene when OC2 is activated because elevated H3K27ac signal was seen across the entire genomic locus only under OC2 OE conditions (Figure 5E). Newly formed enhancer-promoter chromatin loops were seen at the *SRRM4* locus with OC2 OE (Figure 5F). Notably, SEs associated with AR downstream targets in the control cells, including *EHF*, *TMPRSS2*, and *FOLH1* (encoding PSMA), were repressed by OC2 OE (Figure 5E, Figure S5F). Loss of three-dimensional contacts around the *TMPRSS2* promoter were also observed in the OC2 OE condition (Figure 5F-G). These findings confirm that AR is globally suppressed by OC2. The gene set reflecting presumptive OC2-associated, SE-activated or-repressed genes was consistently expressed in the OC2 high/low tumors in the SU2C CRPC cohort (Figure S5G).

In accordance with the robust increase in SRRM4 expression with OC2 OE, SRRM4 activity was similarly increased under these conditions (Figure 5H, Figure S5H). SRRM4 mediates neural-specific alternative splicing [22]. One key downstream SRRM4 target gene is the RE1 silencing transcription factor (REST), a master repressor of neurogenesis. SRRM4-mediated splicing transforms REST into REST4, a transcriptionally inactive form, resulting in loss of the C-terminal repressor domain and diminished repressive function arising from cistrome competition [7]. Elevated REST4 expression was seen in the OC2 OE condition, along with de-repression of a REST-target gene signature (NRSF_01 from MSIGDB [34], and REST translocation into the nucleus was lost (Figure 5I). Depletion of OC2 using siRNA in OC2 OE failed to inhibit the expression of SRRM4 and its targets (Figure S5I), suggesting that the SE formation mediated by OC2 is irreversible. Collectively, these findings imply that OC2 regulates lineage plasticity, at least in part, through SE reprogramming.

### OC2 is a direct suppressor of active androgen

Mass spectrometry analysis of mCRPC tissue [35] showed that OC2 activation determined by gene signature inversely correlated with levels of dihydrotestosterone (DHT), testosterone and androstenedione (Figure 6A). These findings suggest a role for OC2 in androgen metabolism. DHT is inactivated in prostate cells by two glucuronidating enzymes, UGT2B15 and UGT2B17 [36, 37], which suppress AR when activated (Figure 6B). UGT2B15 and UGT2B17 expression was examined in patient tumors [27] with either high or low OC2 activity. Tumors with high OC2 activity exhibited increased levels of UGT2B15 and UGT2B17 overall (Figure 6C). Consistent with this, in LNCaP cells, UGT2B15 and UGT2B17 were upregulated by OC2 OE and suppressed when OC2 was silenced (Figure 6D).

**Fig 6.**
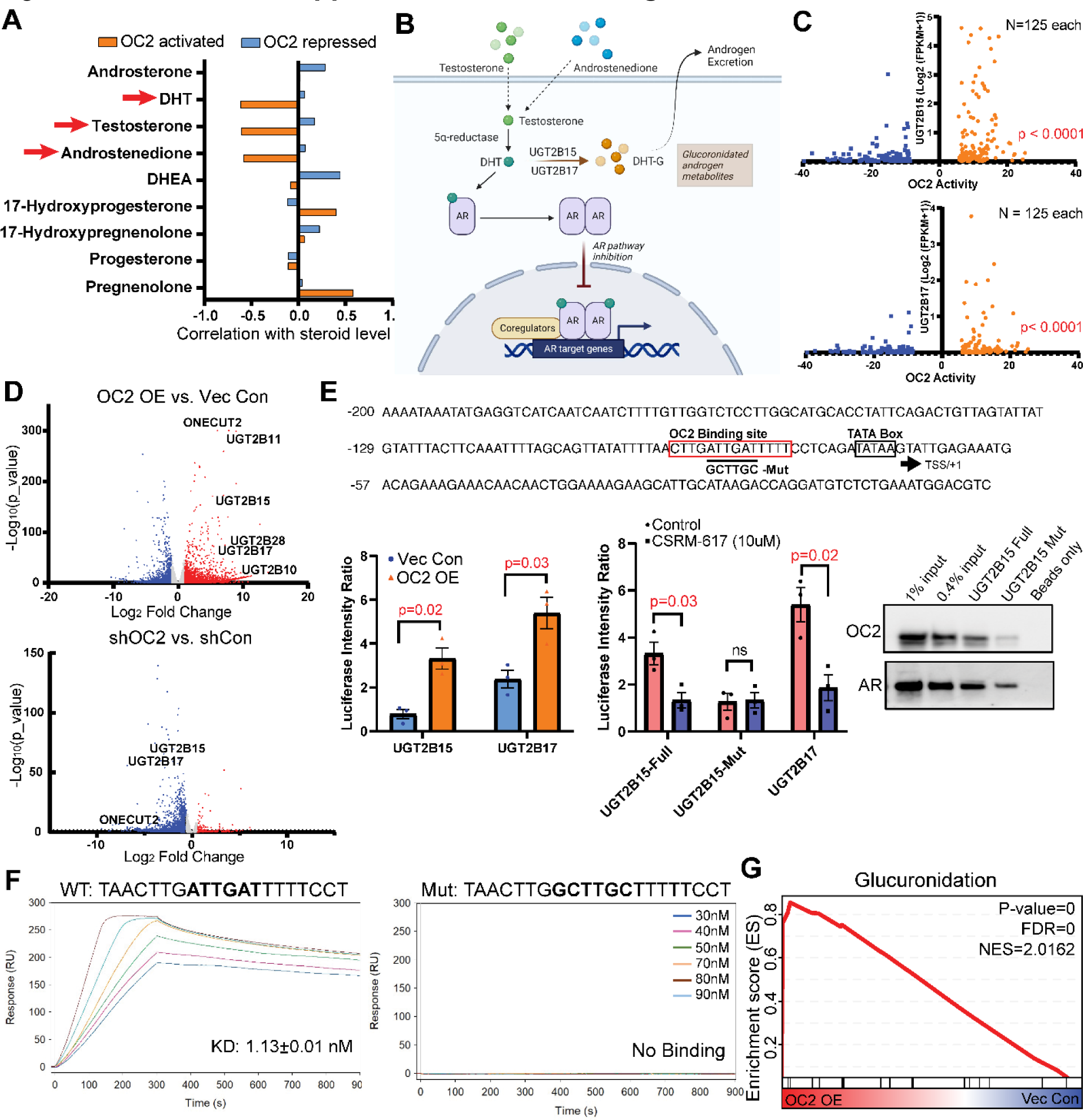
OC2 is a direct suppressor of active androgen. (A) mCRPC patient specimens showing OC2 activity inversely correlated with DHT, testosterone and androstenedione levels. (B) A graphic summary of UGT2B15 and UGT2B17 regulation of androgen glucuronidation to suppress the AR axis. (C) UGT2B15 and UGT2B17 are highly expressed in patients with high OC2 activity. (D) RNA-seq data showing OC2 overexpression induced UGT2B15/17 upregulation while knockdown of OC2 suppressed expression of both genes. (E) Loss of OC2 binding to the *UGT2B15* promoter region when the predicted binding site is mutated. Luciferase reporter system showing OC2 upregulates UGT2B15 and UGT2B17. The luciferase signal with the OC2 binding site mutation was not affected by the OC2 inhibitor. (F) In a cell-free system using surface plasmon resonance, the OC2 DNA binding region exhibited high affinity to the wild-type UGT2B15 promoter while the mutated DNA sequence (90 nM) exhibited no binding signal. (G) Glucuronidation is highly activated in the OC2 OE condition.

OC2 bound to the *UGT2B15* and *UGT2B17* promoters in proximity to AR and FOXA1 (Figure S6A). Motif scanning identified a putative OC2 binding site near the TATA box of *UGT2B15 and UGT2B17* promoters (Figure S6B). UGT2B15 and UGT2B17 promoter-luciferase constructs were significantly activated in OC2 OE vs. control cells (Figure 6E). A 6-bp mutation was introduced at an OC2 site within the *UGT2B15* promoter and incorporated into a promoter (*UGT2B15*-2315/+24)-reporter system (Figure 6E). While the OC2 inhibitor CSRM-617 [4] suppressed wild-type UGT2B15 and UGT2B17 promoter activity, the mutant UGT2B15 promoter showed reduced activity in comparison to wild-type and did not respond to the inhibitor (Figure 6E). Biotinylated (25bp) DNA probes corresponding to the OC2 motif in the *UGT2B15* promoter were synthesized to pull down possible binding proteins from nuclear lysates. OC2 bound to this region, however OC2 binding was lost with the mutation, consistent with this site being an essential element for *UGT2B15* regulation (Figure 6E). AR is a direct repressor of UGT2B15 and UGT2B17 expression [29, 30]. AR bound to both the wild-type and mutant *UGT2B15* 25 bp segment. Of note, the AR inhibitor enzalutamide only activated reporter activity from the wild-type but not the mutant construct, indicating that mutation of the OC2 binding site alters the response of this region to AR suppression (Figure S6C).

OC2 binding to this region was validated using surface plasmon resonance. Recombinant, purified OC2 demonstrated high affinity (K_D_ = 1.13±0.01 nM) to the OC2 DNA binding region in the wild-type *UGT2B15* promoter while showing no binding to the mutated fragment (Figure 6F). Comparison of the 5’ flanking region of several UGT2B genes demonstrated that the OC2-binding region is conserved across several UGT2B family members (Figure S6D). We also observed that enforced OC2 exerted broader effects on activation of glucuronidation genes (Figure 6G). These findings indicate that OC2 is a direct transcriptional activator of androgen-inactivating proteins that irreversibly deplete intracellular androgen. Because a low level of androgen is sufficient to promote lineage plasticity in PC cells [38], these findings identify another mechanism, distinct from those described above, whereby OC2 promotes the emergence of lineage variants.

### OC2 inhibition suppresses a lineage plasticity program induced by enzalutamide

Enzalutamide induces lineage plasticity in AR-dependent PC cells through transcriptional reprogramming [39, 40]. OC2 was previously shown to be inactivated by REST [4]. Enzalutamide-treated LNCaP cells suppressed REST and upregulated OC2 expression in a time-dependent manner (Figure S7A). In two independent patient cohorts, OC2 activation is seen in response to ARSI therapy in pre- and post-enzalutamide matched samples (Figure 7A).

**Fig 7.**
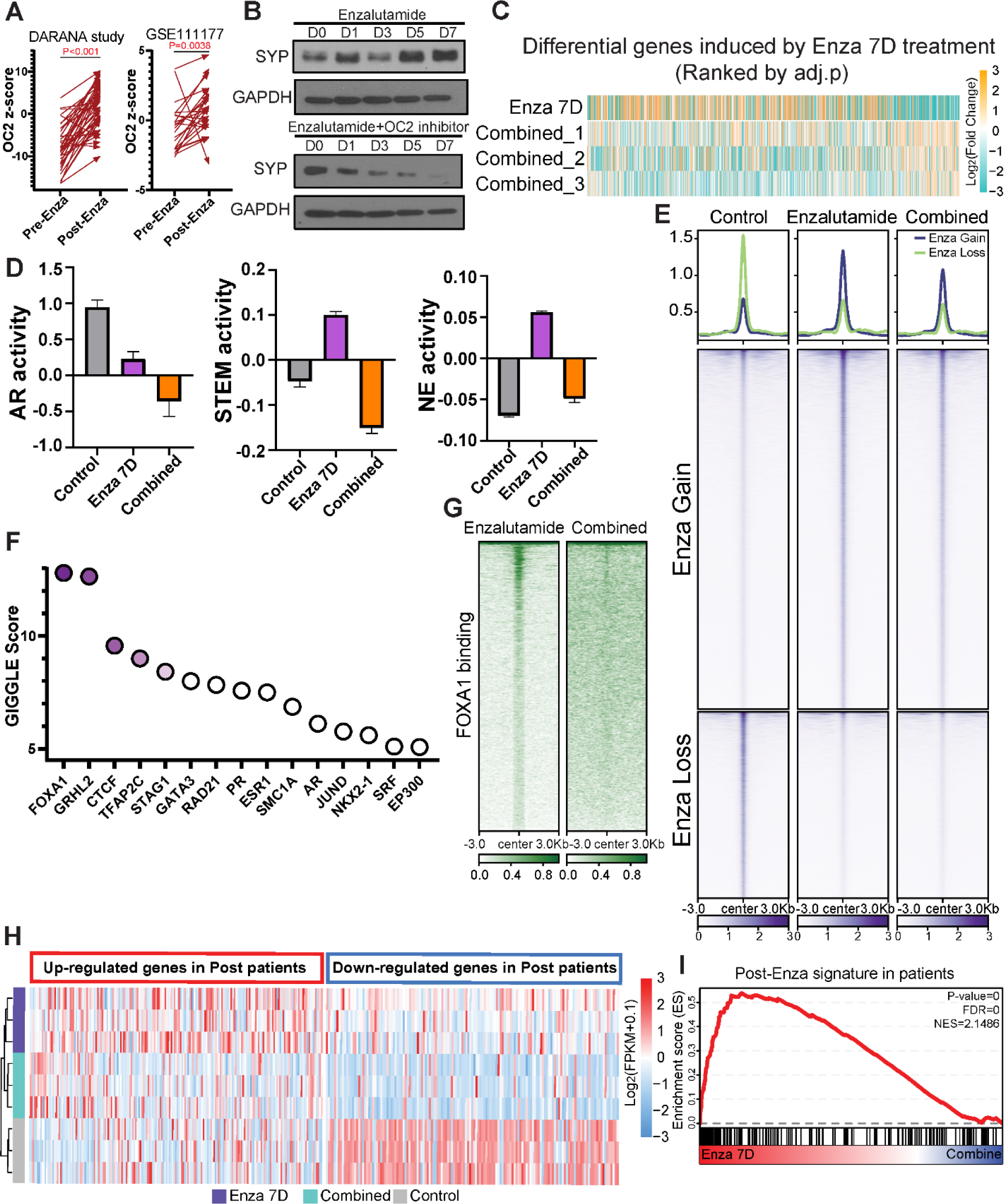
OC2 inhibition suppresses lineage plasticity evoked by enzalutamide. (A) Pre- and post-enzalutamide treatment in the same patients showed OC2 activation following ARSI therapy (GSE197780 GSE111177). (B) OC2 inhibitor blocks enzalutamide-induced SYP expression in LNCaP cells. (C) Simultaneous combination (Combined) treatment of OC2 inhibitor with enzalutamide broadly suppressed enzalutamide-induced gene expression changes vs. control. (D) AR, Stem and NE lineages were repressed by combined treatment with OC2 inhibitor. Heatmap of gene expression in these three lineages is shown. (E) Combined treatment with OC2 inhibitor greatly suppressed enzalutamide-induced chromatin accessibility changes. (F). GIGGLE analysis identified FOXA1 as the most enriched TF in suppressed hyper-accessible regions of combination treatment vs. enzalutamide alone. (G) FOXA1 CUT&RUN-seq showed combined treatment with OC2 inhibitor suppressed FOXA1-driven chromatin accessibility changes. (H-I) A gene signature derived from post-enzalutamide patient samples was consistent with perturbed genes seen in enzalutamide-treated LNCaP cells. SC treatment with OC2 inhibitor blocked enzalutamide-induced gene perturbation.

We applied two protocols to assess the outcomes of combining a direct OC2 inhibitor with enzalutamide: 1) a simultaneous protocol where the both agents were applied to cells at the same time (hereafter referred to as “combined”); and 2) a condition where the OC2 inhibitor was applied alone three days before the simultaneous administration of the both agents (hereafter referred to as “pre-treatment”).

Combined treatment of LNCaP cells with enzalutamide and the OC2 inhibitor CSRM-617 suppressed SYP induction (Figure 7B) and broadly suppressed enzalutamide-induced gene expression changes (Figure 7C; Figure S7B), indicating the potential of targeting OC2 to suppress lineage plasticity (Figure 7B). Combined treatment with OC2 inhibitor robustly suppressed inflammatory response and EMT processes compared to enzalutamide alone (Figure S7C). This finding aligns with a recent study indicating that an inflammatory response is required to induce lineage plasticity in PC cells [16]. OC2 OE significantly released AR binding at *SOX2* and *POU3F2* (BRN2) loci, leading to activation of Stem and NE lineage genes (Figure S7D). Enzalutamide treatment alone activated NE+/Stem+ and NE+/Stem-lineages, while both were suppressed in the combined condition. AR activity was suppressed with enzalutamide, and combined treatment with the OC2 inhibitor further repressed AR (Figure 7D, Figure S7E), consistent with the results shown in Figure S4C.

Enzalutamide used alone induced chromatin remodeling as reported [41] (Figure 7E). However, combined treatment with OC2 inhibitor greatly suppressed enzalutamide-induced chromatin accessibility changes. FOXA1 was identified as the most enriched TF at chromatin regions where OC2 inhibitor suppressed chromatin accessibility, suggesting that OC2 blocks FOXA1-mediated accessibility in enzalutamide-treated cells (Figure S7F, Figure 7F). FOXA1 was shown to be involved in treatment-emergent NE transdifferentiation [42]. FOXA1 binding intensity was dramatically lost with combined treatment, demonstrating that OC2 inhibition blocks enzalutamide-induced lineage plasticity partially by suppressing FOXA1-mediated chromatin remodeling (Figure 7G).

To assess the clinical relevance of the combined treatment strategy, a gene signature derived from post-enzalutamide patient samples was perturbed in enzalutamide-treated LNCaP cells consistent with the patient data. Combined treatment blocked this enzalutamide-induced gene perturbation. (Figure 7H and I). These findings demonstrate that OC2 inhibition suppresses an enzalutamide-induced gene expression network seen in human PC.

The use of OC2 inhibitor alone activated AR signaling (Figure S7G), suggesting that OC2 inhibition was capable of sensitizing cells to enzalutamide. To test this, a second treatment protocol was used. LNCaP cells were pretreated with CSRM-617 for three days, followed by combined treatment with enzalutamide. This sequential “pre-treatment” protocol was extremely potent in tumor cell killing compared to the combined condition (Figure S7H). RNA expression profiling of the pre-treatment condition revealed predominant effects on cell cycle processes in contrast to the combined protocol, consistent with the cell growth and survival results, suggesting that a therapeutic window exists for OC2 targeting even before ARSI therapy (Figure S7I).

## DISCUSSION

OC2 was identified previously as an NE driver that can be inhibited directly with a small molecule capable of causing regression of established human mCRPC metastases in mice [4]. In the present study, we characterized the mechanisms whereby OC2 drives lineage plasticity. We found OC2 to be widely expressed in CRPC tumors, and that it functions as a broadly acting lineage facilitator through its ability to alter chromatin accessibility, promote the formation of SE regions, and regulate gene expression by acting at bivalent (poised) promoters. OC2 suppresses the AR through multiple mechanisms, including promoting androgen inactivation through activation of glucuronidation genes that specifically and irreversibly disable androgens. We further show that these diverse mechanisms promoting plasticity include direct activation of *NR3C1*, the gene encoding the GR. The GR promotes disease progression in PC by assuming direct control of part of the AR cistrome under conditions of AR suppression, thereby constituting a mechanism of resistance to AR-targeted therapy. Notably, activation of the GR occurs in adenocarcinoma, not NEPC. We show here that, through the intercession of the GR, OC2 can activate certain AR-regulated genes. Consequently, the ability of OC2 to promote lineage variation extends beyond NEPC to treatment-resistant adenocarcinoma.

We present evidence that OC2 directly activates multiple AR-independent, lineage-defining factors, including NR3C1, ETV4, TWIST1, BRN2 (POU3F2), KLF5 and TFAP2A. ETV4 mediates dosage-dependent prostate tumor initiation and promotes metastasis of prostate adenocarcinoma in response to PI3K and RAS co-activation [43]. KLF5 opposes AR activities and drives the double-negative PC phenotype [26, 44]. TFAP2A activation indicates emergence of neural crest lineages [25]. In contrast, classical AR downstream target genes (*EHF*, *TMPRSS2*, *KLK3*) were suppressed by OC2. Beyond direct epigenetic regulation within promoter regions, gene expression changes were also contributed by reprogrammed enhancers looped to these promoters. Consistent with the conclusion that OC2 has the capability to bind to closed chromatin and promote chromatin accessibility, we show that OC2 interacts with SMARCA4 and SMARCA5, which are important components of CRCs that modify chromatin architecture to allow TF access to condensed chromatin [45, 46]. Inactivation of CHD1, another chromatin remodeler, has been shown to promote the emergence of tumor heterogeneity and enzalutamide resistance [47]. Notably, all four lineages driven by CHD1 loss (GR-driven, BRN2-driven, NR2F1-driven and TBX2-driven EMT) emerge with OC2 activation. Taken together, these results indicate that OC2 facilitates the development of multiple lineages through chromatin remodeling and epigenetic modification.

We also demonstrate that OC2 upregulates the *UGT2B15* and *UGT2B17* genes and is a direct transcriptional activator of *UGT2B15*. The UGT2B15/17 proteins irreversibly inactivate T and DHT. Analysis of human CRPC tumors revealed that tumors with high levels of OC2 activation exhibited reduced levels of T, DHT, and androstenedione. These findings indicate that OC2 promotes clearance of androgen, thus ensuring a low androgen environment. Reduced levels of androgen promote AR-indifferent lineage variation in PC cells [38]. Consequently, OC2 can also facilitate lineage variation by activating gene expression programs suppressed by the AR. RNA expression in PC models, and analysis of human PC cohorts, indicate that OC2 exerts a broad effect on glucuronidation processes generally. Upregulation of glucuronidation has been reported to induce multi-drug resistance [48].

Treatment of PC with the clinical ARSI, enzalutamide, evokes a gene expression program associated with lineage plasticity [39, 40]. We validated this finding here, and we further demonstrated that enzalutamide activates OC2. Simultaneous treatment with OC2 inhibitor and enzalutamide blocked enzalutamide-induced gene expression changes and suppressed enzalutamide-induced chromatin accessibility, suggesting that OC2 inhibition may suppress lineage infidelity evoked by hormone suppression.

In summary, our results identify OC2 as a novel facilitator of multiple AR-indifferent lineages, operating by parallel routes to support the appearance of treatment-resistant adenocarcinoma as well as NEPC variants. Our work suggests that OC2 inhibition could play an important role in blocking or delaying the emergence of CRPC.

### Potential Conflicts

P.S. Nelson served as a paid advisor for Bristol Myers Squibb, Pfizer, and Janssen and received research support from Janssen for work unrelated to the present study.

## Acknowledgments

This work was supported by UCLA Prostate Cancer SPORE 2P50CA092131, 1R01CA220327, 1R01CA271750; Department of Defense PC180541, PC190604, PC190482; The Institute for Prostate Cancer Research, the Pacific Northwest Prostate Cancer SPORE P50CA97186, R01CA266452, and R01 CA234715. The authors thank Dr. Gerhard Coetzee for helpful discussions.

## METHODS

### Data availability

We used a series of global data publicly available and generated new global data for this study (**Table S1**). Initially, we obtained a digital spatial profiling (DSP) transcriptome data from Gene Expression Omnibus (GEO) database (GSE147250) [6]. Single-cell transcriptome from PC patients, were integrated with three independent data sets (GSE141445, GSE157703, and GSE137829) [9–11], consisting of 6 CRPC and 14 primary samples. We also included tissue transcriptome data (GSE147493) of non-metastatic (M0) and metastatic (M1) PC needle biopsy samples from Great Los Angeles VA cohort [17] and the data of PC samples from various metastases sites in Stand-Up-To-Cancer (SU2C) cohort [27], which was downloaded from cBioPortal [49]. The processed PC transcriptome data (level-3) from The Cancer Genome Atlas (TCGA) collection were downloaded from Genomics Data Common (released on March 26, 2019; GDC V16.0). The global histone mark profiles, includingH3K27ac, H3K4me3 and H3K27me3 from LuCaP PC patient-derived xenograft (PDX) [42] were included for this study. Finally, drug-induced tissue transcriptome data from PC patients by enzalutamide treatment were obtained with the accession, GSE197780 [50]. All global data generated for this study were deposited to GEO database with accessions (GSE244025).

### Gene signature collection

We make a collection of gene signatures for this study based on the previously published literatures and Molecular Signature Database (MSigDB) [8]. The ‘Hallmark Androgen Response” gene set as an AR signature was from MSigDB. NE and stem signatures were collected from published literatures [7, 51]. All the literatures and gene list for each signature were included in **Table S2**.

### Digital spatial profiling (DSP) data analysis

The 152 cores from 53 tumors were ranked by OC2 *in situ* hybridization levels from high to low. Immunohistochemical profiles of PSMA and FAP proteins AR and NE signature scores were also displayed at the corresponding tumor cores. Known DSP class information of the 53 tumors also labeled in Figure 1A [6]. The signature scores were computed by using z-score method [52] with AR, NE and stem signature genes in each tumor core. The scores and expression levels of the 152 cores were visualized using heatmap.

### Single cell transcriptome data analysis

#### Preprocessing and data integration

We found three scRNA-seq data form previous studies [9–11]. For each data, doublets and triplets were removed using DoubletFinder [53]. By default, 2,000 highly variable genes were selected in each dataset, which was used in Principal Component Analysis (PCA). Then, the cells with the number of genes < 500 and >20% mitochondrial expression were removed (nFeature_RNA > 500 & percent.mi < 20%), resulting a total of 36,419 primary and 23,607 CRPC cells. Systemic biases from the individual data were adjusted through the canonical correlation analysis with Seurat R package (version 4.3.0) [54].Batch effect was removed in data combination.

#### Dimensionality reduction

30 principal components were selected based on statistical significance from Elbow plot analysis. We then performed the Uniform Manifold Approximation and Projection (UMAP) analysis with the 30 selected principal components to identify cell sub-populations (arXiv:1802.03426). Clustering was performed for integrated expression values using the Louvain community detection-based method with *FindNeighbors* function in Seurat package with the resolution is 0.8. To visualize the cell clusters, UMAP plot was used. The following markers were used for cell type annotation: Epithelial/Tumor: EPCAM, KRT8; T cells: CD3E, CD3D, TRBC1/2, TRAC; Myeloid cells: LYZ, CD86, CD68, FCGR3A; B cells and plasma cells: CD79A/B, JCHAIN, IGKC, IGHG3; Endothelial cells: CLDN5, FLT1, CDH1, RAMP2; Fibroblasts: DCN, C1R, COL1A1, ACTA2; Smooth muscle cells: TAGLN, CNN1; Mast cells: TPSAB1, resulting 8 cell types consisting of 36,419 primary and 23,607 CRPC cells were identified (**Figure S1E**). In order for in-depth analysis of OC2 expressing cells, we selected OC2 expression cells with positive gene expression level (Figure 1C). We then computed weighted z-score of the gene sets, including NE markers (CHGA, ASCL1 and BRN2), Androgen response genes, Stemness signature, and other PC associated gene signatures for each cell type. The activity of top 20 TFs (ranked by average TF activity in each module) were showed on **Figure 1F** using pheatmap package.

#### Single cell trajectory analysis

Pseudotime inference was performed to track the progression trajectory of OC2-expressing cancer cells by monocle3 R package [55]. Epithelial cells from M0 stage in primary samples were selected as the start nodes. The learn_graph function was used to infer the trajectory of all OC2-expressed M0 and CRPC cells. Then, the plot cells function was used to visualize the trajectory and pseudotime inference results. The signature score of each cell was defined by calculating z score.

### Transcription factor (TF) activity inference and lineage identification

The activity level of transcription factor (TF) in a given sample or cell was computed using decouplR package [15]. VIPER method [56] was applied to infer TF activity based on mRNA expression of its targets including all targets with confidence levels from A to D [14]. We identified cell type clusters defined by the TF activities using K-means method in ConsensusClustering package [57]. Considering the known CRPC lineages and the Consensus Cumulative Distribution plot, the optimal number of clusters (K) was identified by surveying K values from 2 to 8. This was visualized using heatmap with TF activities of the cells in the 4 clusters, in conjunction with AR, NE and stemness gene signature scores.

### Cell lines

LNCaP (#CRL-1740) was obtained from the American Type Culture Collection (ATCC) and authenticated using the Promega PowerPlex 16 system DNA typing (Laragen). LNCaP cells were grown in RPMI-1640 media (Gibco) supplemented with 10% FBS and penicillin/streptomycin. LAPC4 cells was gifted from Dr. Stephen J. Freedland group. LAPC4 cells were grown in IMDM (Gibco) supplemented with 10% FBS and 1nM R1881 plus penicillin/streptomycin. Mycoplasma contamination was routinely monitored using the MycoAlert PLUS Mycoplasma Detection Kit (Lonza #LT07-118). The OC2 overexpression construct was generated by cloning the full-length OC2 cDNA (NM_004852) into the pLenti-C-Myc-DDK-IRES-Puro (Origene #PS10069) lenti-virus system. The inducible OC2 was generated with pCDH-CuO-MCS-IRES-GFP (SystemBio #QM530A-2) and pCDH-EF1-CymR-T2A-RFP (SystemBio #QM300PA) system with Cumate (SystemBio # QM100A) turned-on treatment. The AR-respond plasmid ARR3tk-eGFP/SV40-mCherry was purchased from Addgene (Addgene, #132360). Then packing (psPAX2, Addgene #12260), and envelope (pMD2.G, Addgene #12259) plasmids were co-transfected into HEK293T cells to produce lentivirus. Cells were infected with lentivirus supplemented with 10 µg/mL polybrene, then selected by 2ug/mL puromycin to generate the stable overexpression cells.

### SCID mouse *in vivo* xenografts

All experimental protocols and procedures were approved by the Institutional Animal Care and Use Committee (IACUC) at Cedars-Sinai Medical Center. All relevant ethical regulations, standards, and norms were rigorously adhered to. For In vivo tumor growth assays, LNCaP control or OC2 OE cells were adjusted to 1 × 10^7^ cells/mL in DPBS, followed by mixing with Matrigel at a 1:1 ratio (v/v). For each male SCID/Beige mouse (7-weeks-old; Charles River #CRL:250; CB17.Cg-PrkdcscidLystbg-J/Crl), 100 µL mixture was subcutaneously injected into both flanks. Tumor length and width were measured with a caliper and tumor volumes were calculated using the formula of (length × width2)/2. At the endpoint, mice were euthanized, and tumor xenografts were collected and send to Cedars Sinai Pathology Laboratory for preparing FFPE slides.

### Tissue Sample Preparation

Formalin-fixed paraffin-embedded (FFPE) slides were obtained from TMA 95A, B, and C, provided by the University of Washington collected from CRPC rapid autopsy patients who signed written informed consent under the aegis of the Prostate Cancer Donor Program at the University of Washington (IRB protocol no. 2341), to which the sections cut from all the positive and negative tissues and cell blocks were added on each slide. The positive controls included fetal Retina, Pancreas, Xenograft OE2, and HistoGel pre-wrapped cells block of organoid BS17077 with force expressed OENCUT2. The negative control consisted of organoid BS17077 with no vehicle and Xenograft Vec-Con1.

### *In Situ* Hybridization

For in situ hybridization, the RNAscope® 2.5 HD detection reagent–Red kit (ACD, 322360) was utilized according to the manufacturer’s instructions. The probes used in this study included the Hs-ONECUT2 probe (ACD, 473531) targeting the ONECUT2 mRNA, the probe_DapB (ACD, 310043) as the negative control, and the Probe_PPiB (ACD, 313901) for the positive control.

### Immunohistochemistry

The FFPE slides were deparaffinized, and antigens were unmasked using EnVision FLEX Target Retrieval Solution Low PH in DAKO PT link at 99°C for 20 minutes. Subsequently, the slides were loaded onto the Agilent Automated AS48 Link. Background reduction was achieved with EnVision Flex Peroxidase and Casein blocking agents. The primary antibodies used in this study included rabbit anti-human monoclonal antibodies of ONECUT2 (Sigma HPA057058) at 1:50, Androgen Receptor (Cell Marque, SP107) at 1:6400, Glucocorticoid Receptor (D6H2L) XP® (Cell Signaling, 1204) at 1:6000, and Synaptophysin (D8F6H) XP® at 1:200. Visualization was carried out with EnVision Flex HRP labeled Polymer (Agilent SM802), followed by development with Flex DAB+ sub-chromogen and counterstaining with EnVision Flex hematoxylin. A peptide challenging assay was conducted using PrEst Antigen ONECUT2 (Sigma APrEST86051) with a 5:1 ratio of peptide to ONECUT2 antibody, pre-incubated for 1 hour at room temperature, Flex mouse (cocktail IgG1, 2a,2b,3 and IgM) and Immunoglobin fraction of serum from non-immunized rabbits to replace primary antibodies, as well as omitting the primary antibodies, were run with the same protocol parallel with experimental slides as negative controls.

### Western blot analysis

Cell lysates were separated on 4–20% SDS-PAGE (Bio-Rad Laboratories) and transferred to nitrocellulose or PVDF membranes. The membranes were blocked with 5% non-fat dry milk and subsequently incubated with the pertinent primary antibody overnight. Anti-Glucocorticoid Receptor (D6H2L) (CST, #12041, 1:1000 dilution), Anti-AR (Activemotif, #39781, 1:2000 dilution), Anti-OC2 (Proteintech, #21916-1-AP, 1:1000 dilution), Anti-PSA (CST, #5365, 1:2000 dilution), Anti-Histone-H3 (CST, #9715, 1:2500 dilution), Anti-SMARCA4 (Abcam, #ab110641, 1:10000 dilution), Anti-SMARCA5 (Abcam, #ab72499, 1:2500 dilution), Anti-Myc-tag (CST, #2276, 1:1000 dilution), Anti-SYP (CST, #36406, 1:1000 dilution), Anti-REST (Sigma-Aldrich, #ZRB1455, 1:1000 dilution). Membranes were subsequently washed with TBST (0.1% Tween-20) and incubated with an HRP-conjugated secondary antibody (GE Healthcare Bio-Sciences). After washing with TBST, the protein bands were detected by the Chemidoc MP imaging system (Bio-rad).

### Cell proliferation analysis

All procedures were performed according to the XTT cell viability kit protocol (CST, #9095). To assay viability, cells were plated at a density of 2,000 cells/well in triplicate. 48 hours after indicated treatment, viability was assessed at the absorbance of 450nM. IC_50_ was generated by a non-linear regression function in Graphpad 9.0. For proliferation assay, the cells were seeded at 2000/well and treated up to 96 hours, then 450nM absorbance was collected for further analysis.

### RNA extraction and quantitative real-time polymerase chain reaction (RT-PCR)

Total RNA was extracted from cells using Qiagen RNeasy Kit (Qiagen #74104). Messenger RNA was converted to the first-strand cDNA using iScript cDNA synthesis kit (Bio-Rad #1708891), followed by RT-PCR reaction using PowerUp SYBR Green PCR Master Kit (Applied Biosystems #A25742) in QuantStudio 5 Systems (Applied Biosystems #4309155). Gene expression was normalized to Actin/GAPDH using the comparative CT method.

### Interactome Analysis by Immunoprecipitation-Mass Spectrometry (IP-MS)

ChromoTek iST Myc-Trap Kit (Proteintech, #ytak-iST) was used for pulling down interactome with OC2-myc tagged protein. Eluted proteins were analyzed by gel-enhanced liquid chromatography-tandem mass spectrometry (GeLC-MS/MS) essentially as described [58]. The resulting tryptic peptides in 10 μL solution was loaded onto a 2-cm trap column (Thermo Scientific) and separated on a 50-cm EASY-Spray analytical column (Thermo Scientific) heated to 55°C, using a 1-h gradient at the flow rate of 250 nL/min. The resolved peptides were ionized by an EASY-Spray ion source (Thermo Scientific), and mass spectra were acquired in a data-dependent manner (DDA) in an Orbitrap Fusion Lumos mass spectrometer (Thermo Scientific). MS1 scans were acquired in 240,000 resolution at *m*/*z* of 400 Th, with a maximum injection time of 250 milliseconds and an ion packet setting of 4 × 10^5^ for automatic gain control (AGC). Most intense peptide ions with charge state of 2-7 were automatically selected for MS/MS fragmentation by higher energy collisional dissociation, using 30% normalized collision energy. MS/MS spectra were acquired in the ion trap, using rapid ion trap scan at 1 × 10^4^ AGC and 35 millisecond maximum injection time. Dynamic exclusion was enabled to minimize redundant MS2 acquisition.

The acquired RAW files were searched against the Uniprot_Human database (released on 03/30/2018, containing 93,316 protein sequences) with MaxQuant (v1.5.5.1) [59]. The searching parameters include trypsin/P as the protease; carbamidomethyl (C) as fixed modification; oxidation (M), acetyl (protein N-term), and deamidation (NQ) as variable modifications; minimal peptide length as 7; up to two missed cleavages; mass tolerance for MS was 4.5 ppm and for MS/MS was 0.5 Da; identification of second peptides enabled; label free quantification (LFQ) enabled, with match-between-runs within 0.7 min. A standard false discovery rate of 0.01 was used to filter peptide-spectrum matches, peptide identifications, and protein identifications.

### RNA-sequencing and data processing

RNA concentration, purity, and integrity were assessed by NanoDrop (Thermo Fisher Scientific Inc.) and Agilent Bioanalyzer. 1ug Total RNA were shipped to BGI Americas for Pair-end 100bp sequencing using Strand Specific Transcriptome Library Construction Protocol and DNBSEQ platform. FastQC was applied to analyze Raw RNA-Seq fastq files. Then, Trim galore was used to remove the adapters. 150 bp paried-end reads were aligned to human reference genome (HG38) using STAR (-alignIntronMin 20-alignIntronMax 1000000 - alignSJoverhangMin 8-quantMode GeneCounts) method [60]. Then, gene read counts matrix were used for further analysis. Differentially expressed genes were determined using edgeR packages [61]. GSEA Preranked function was performed to identify significant biological functions [62]. Genes were ranked by the log2 fold change between control and OC2 perturbed samples. ClusterProfiler R package was used to perform GO and KEGG pathway enrichment analysis based on differentially expressed gene set [63].

### Cleavage under targets and release using nuclease (CUT & RUN) sequencing

All procedures were performed according to the manufacturer’s protocol from cell signaling technology (CST, #86652). Briefly, 100,000 cells from both control and OC2 OE cells were resuspended in wash buffer (20 mM HEPES-NaOH pH 7.5, 150 mM NaCl, 0.5 mM spermidine, and protease inhibitor cocktail), concanavalin A-magnetic beads added, then rotated for 10 min at room temperature. Cell-bead conjugates were resuspended in 200 µL of digitonin buffer (wash buffer with 2.5% digitonin solution) containing 2ug of Anti-Tri-Methyl-Histone H3 (Lys27) (CST, #9733), 2ug of Anti-Tri-Methyl-Histone H3 (Lys4) (CST, #9751), 2ug of Anti-Acetyl-Histone H3 (Lys27) (D5E4) (CST, #8173), 2ug of Anti-Mono-Methyl-Histone H3 (Lys4) (D1A9) (CST, #5326), 5ug of Anti-ONECUT2 (Proteintech, #21916-1-AP), 5ug of Anti-Glucocorticoid Receptor (D6H2L) (CST, #12041), 5ug of Anti-AR (Activemotif, #39781), 5ug of Anti-FOXA1 (Activemotif, #398837) primary antibody, or rabbit IgG (CST, # 66362,), rotated overnight at 4 °C, resuspended in 250 µL of antibody buffer and 7.5 µL of the pAG-MNase enzyme (# 57813, CST), followed by the rotation at 4 °C for 1 h. After washing with digitonin buffer, ice-cold 150 µL of digitonin buffer containing CaCl2 was added and incubated on ice for 30 min followed by the addition of 150 µL of stop buffer containing 5 ng *S. cerevisiae* spike-in DNA used for sample normalization. After incubation at 37 °C for 15 min, samples were centrifuged at 16,000 g for 2 min at 4 °C. Tubes were placed on a magnetic rack, then supernatants were collected. DNA was purified using DNA purification buffers and spin columns (CST, #14209). The CUT & RUN library was generated with the DNA Library Prep Kit for Illumina (CST, #56795) combined with Multiplex Oligos for Illumina® (Dual Index Primers) (CST, # 47538). The adaptor was diluted 1:25 to avoid contamination. The PCR enrichment step run 15 cycles to amplify the adaptor-ligated CUT&RUN DNA.

### CUT&RUN-seq data processing and downstream analysis

CUT&RUN-seq data of H3K27ac, H3K27me3, H3K4me3, H3K4me1 and several TFs (ONECUT2, AR, FOXA1 and GR) was generated in LNCaP cell lines. Briefly, Trim Galore was utilized to remove contaminant adapters and read quality trimming. 150 bp paired-end reads were aligned to human reference genome (HG38) using Bowtie2 (v2.2.6) [64]. As spike-in is commonly used as a control probe in DNA sequencing [65]. We added the *S. cerevisiae* genome as spike-in control reference. Then, the mapping rates were used for calculating scale factor in each sample. Next, Picard MarkDuplicates tool was utilized to mark and remove PCR duplicates in each sample. ENCODE blacklisted regions on HG38 [66] were removed by using bedtools [67]. Finally, a high accuracy peak calling method, SEACR [68], was used to identify significant peaks with the parameters: *norm stringent*. To visualize the signal in each sample, bamCompare function in DeepTools (v3.1.3) was used to generate Bigwig files with the parameters: *-binSize 10-numberOfProcessors 5-normalizeUsing CPM-ignoreDuplicates - extendReads 200* [69]. The scale factor was calculated from the above spike in step. Integrative Genomics Viewer (IGV) was used to visualize the signal in bigwig files [70]. We employed the GIGGLE method [71] to identify the TFs whose genome-wide binding profiles publicly available and produce by this study are highly enriched in the specific peaks.

### Omni-ATAC sequencing

50,000 viable LNCaP Vec_Con and OC2 overexpressing cells were precipitated and kept on ice and subsequently resuspended ATAC Resuspension Buffer (RSB) (49.25mL nuclease-free water, 500uL 1M PH7.5 Tris-HCL, 100uL 5M NaCl, and 150uL 1M MgCl2). 50uL of Transposition Master mix (25uL 2xTD buffer, 1uL Tagment DNA enzyme, 16.5uL of PBS, 0.5uL 1% Digitonin, 0.5uL 10% Tween-20 and 6.5uL Nuclease-free water) for 30min reaction at 37C with 1000 rpm mixing. DNA was then purified using Zymo DNA clean and Concentrator-5 kit. ATAC-seq Libraries were prepared following the Buenrostro protocol [72]. The Tagment DNA enzyme was gifted from Pattenden, Samantha Lab from UNC.

### ATAC-seq data analysis

Trim Galore and Bowtie2 (v2.2.6) were utilized to do read quality trimming and mapping to HG38 reference genome. PCR duplicates and blacklisted regions (HG38) were removed by the same methods described above. After that, Macs2 method was utilized to do peak calling based on the following parameters: *-bdg - SPMR-nomodel-extsize 200-q 0.01*. Bigwig files were also generated by bamCompare function in Deeptools (v3.1.3) using the parameters: *-binSize 10 - numberOfProcessors 5-scaleFactorsMethod None-normalizeUsing CPM - ignoreDuplicates-extendReads 200*. Then, IGV software was used to do visualization. The HOMER software with findMotifsGenome.pl function was used to identify enriched motifs in selected regions. The parameters in findMotifsGenome.pl are hg38-size 200-len 8, 10, 12 [73]. For visualization of the peak profiles from ATAC-seq and CUT&RUN-seq data, we used deepTools (v2.5.0) [69] to generate read abundance from all datasets around peak center (± 3 kb/ 5 kb), using ‘computeMatrix’. These matrices were then used to create heatmaps and profiles, using deep-Tools commands ‘plotHeatmap’ or ‘plotProfile’ respectively. For genome browser track, we used pyGenometracks package [74] to generate plots for tracks visualization (Reference Genome, HG38).

### Mutation constructs UGT2B15 promoter luciferase

Quick Change II XL site-directed mutagenesis kit (Agilent) manufacturers protocol was used to generate mutation constructs. PCR amplification of the UGT2B15 mutation constructs were cycled at 1 cycle 95°C 1 minute, 18 cycles at 95°C 50 seconds 60°C 50 seconds 68°C 1 minute/kb of plasmid length and 1 cycle at 68°C 7 minutes. The PCR product was then then placed on ice for 2 minutes to cool the reactions to ≤37°C. Then enzyme digestion of the PCR product was with the addition of 1 µl of the Dpn I restriction enzyme (10 U/µl).

UGT2B15mut_F_gttgtttctttctgtcatttctcatacttatatctgaggaaaagcaagccaagttaaaatataactgctaaaatttgaagtaaatacataata

UGT2B15mut_R_tattatgtatttacttcaaattttagcagttatattttaacttggcttgcttttcctcagatataagtatgagaaatgacagaaagaaacaac

### Luciferase reporter assay

Cells were co-transfected with PGL4.10 luciferase backbone construct, and the pRL-SV40 vector (Promega #E2231) with Turbofectin 8.0 (Origene, #TF81001). After transfection overnight, cells were washed and treated for 6 hours with CSRM-617 (10uM). Luciferase activity was measured using the Dual-Luciferase Assay System (Promega, #1910).

### DNA-Protein Affinity Assay (DAPA)

10M Cells were harvested and nuclear protein and frozen at −80°C for each DAPA reaction. SRRM4 300bp promoter region was amplified from gDNA with the primer (5’ TTTCTCCTCCCAAGACCTGCG, 5’ TCTGAGCTGGCTGAGCCTCT) using Phusion High-Fidelity PCR Kit (Thermoscientific #F553L). Then the DNA was biotinylated with BioNick DNA Labeling Kit (Invitrogen #18247015). Biotinylated 25bp UGT2B15 WT and Mut probes were synthesized by IDT. Primers were diluted to 20 μM as working stocks for the DAPA reaction. Each primer was incubated with 20 μL of each primer (F/R) incubated in 60 μL of DAPA buffer (10 mM Tris Buffer, pH 7.4, 1 mM EDTA). The primer mix was incubated at 95° C for 3 minutes. The annealed primers were then cooled at room temperature overnight and stored at −20°C. The nuclear protein fraction was isolated using the cytoplasmic and nuclear extraction protocol (Thermofisher # 78833). An aliquot of 250 μg of nuclear protein lysate was diluted in DAPA wash buffer (20 mM Hepes buffer, pH 7.4, 1 mM DTT, 0.1% Tween-20) up to a volume of 500 μL. The nuclear protein lysate then was pre-cleared by adding 2 μL of poly D[IC] (Sigma-Aldrich) 25 µg/µL and 20 μL of High Capacity Strepavidin agarose beads (Thermofisher #20357). The pre-cleared protein lysate was then incubated for 2 hour at 4° C on an end-over-end rotator. The mixture was then centrifuged at 3000xg at 4° C, and the protein lysate in the supernatant fraction was used for binding. The beads and buffer were then boiled at 95°C for 5 minutes and used as a negative control sample. The pre-cleared lysate was bound to 250 ng of the biotynlated primer and an additional 2 μL of poly D[IC]. The protein-DNA mixture was then incubated overnight at 4° C on an end-over-end rotator. The protein-DNA complex mixture was then incubated with 20 μL of High Capacity Strepavidin agarose for 2 hours at 4° C. The mixture was then centrifuged a 3000xg for 1 minute and the supernatant was removed. The pelleted beads were then washed with three times with 500 μL DAPA wash buffer. All wash buffer was then removed and then added 30 µL 4X lammeli buffer supplemented with 5% β-meracptoethanol. The beads and buffer were then boiled at 95°C for 5 minutes then analyzed by western blot.

UGT2B15 WT probe(F) Biotynlated-TTTTAACTTGATTGATTTTTCCTCA UGT2B15 WT probe (R)-TGAGGAAAAATCAATCAAGTTAAAA

UGT2B15 Mut probe (F) Biotynlated-TTTTAACTTGGCTTGCTTTTCCTCA UGT2B15 Mut probe R)-TGAGGAAAAGCAAGCCAAGTTAAAA

### Surface plasmon resonance binding studies

Measurement of OC2 binding affinity for the UGT2B15 promoter DNA sequence was analyzed using surface plasmon resonance with Sartorius Octet SF3 instrument. The UGT2B15 promoter 20 base-pair sequence (5’-TAACTTGATTGATTTTTCCT-3’ for wild type and 5’-TAACTTGGCTGTCTTTTCCT-3’ for mutant) was immobilized to a streptavidin SADH sensor chip (Sartorius) in running buffer (10 mM HEPES pH 6.8, 250 mM NaCl, 0.005% Tween 20). The 20 base-pair reverse complement sequence (100 ng/ml), biotinylated at the 5’ end, was first immobilized to 75 response units (RU), followed by immobilization of the 20 base-pair forward sequence at 100 ng/ml to a final 150 RU, ensuring duplexed DNA immobilization. Recombinant OC2 DNA-binding domain protein (L330-W485), purified as previously published (Nat Med ref), was diluted in running buffer to 30, 40, 50, 70, 80, and 90 nM and injected over the immobilized, duplexed 20 base-pair DNA probe as well as reference channel. In the case of mutant UGT2B15, where no binding was observed, a subsequent positive control of wild-type UGT2B15 was performed on the same channel to confirm adequate experimental setup. The sensograms were fit with Octet SPR Analysis software (Sartorius) using a one binding site model with mass transport correction to calculate binding affinity (KD). All DNA oligos were purchased from IDT (synthesized in San Diego, CA, USA).

### Genome-wide chromatin interaction analysis using Hi-C

Hi-C data was generated using the Arima-HiC kit, according to the manufacturer’s protocols [19, 20]. The Juicer pipeline (version 1.6) was run on all samples using default parameters [75]. The reads were aligned to the hg38 reference genome. Chromatin loop calls were identified using HiCCUPS (Hi-C Computational Unbiased Peak Search) algorithm with KR (Knight-Ruiz) normalization. TADs (Topologically Associating Domains) were identified using Arrowhead algorithm at 10kb resolution. Eigenvalues were identified using Eigenvector algorithm at 1Mb resolution across the genome. Then multiple .txt files containing eigenvalues were converted and merged into a single .wig file containing final A/B compartments calls.

### ATAC-Seq and RNA-Seq data integration

To integrate gene expression (RNA-seq) and chromatin accessibility profiles (ATAC-seq) induced by OC2, we first performed differential expression analysis using edgeR, and differential ATAC-seq peaks using the Differential ATAC-seq Toolkit (DAStk) [76]between control and OC2 OE. We then computed the z-scores for differential expression and differential motif enrichment for each TF based on the testing P-values from edgeR and DAStk tools, respectively. The z-scores were combined into an overall z-score using the Stouffer’s Method [77] as the TF activity score. Finally, the significance level of the TF activity score was calculated by converting the combined z-score to P-value with the standard normal cumulative distribution function. The positive and negative z-score represent the active and repressed status of TFs by the OC2 perturbation. For further consideration of TF activity. We estimated the downstream target gene contribution to TF activity by calculating the Fold Enrichment Score based on the TF-target interaction information from Dorothea R packages [78]. The TF fold enrichment was defined as the percentage of the number of differentially expressed target genes in the list belonging to all the TF target genes, divided by the corresponding percentage of the number of DEGs in all genes in the data. The higher Fold Enrichment Score is the more downstream target genes changed at expression level. The significant TFs were selected with p-value<0.05 and Fold Enrichment Score≥1.5.

### Promoter H3K4 and H3K27 trimethylation analysis

Counts per million (CPM) mapped reads value from H3K4me3 and H3K27me3 CUT&RUN-seq data within ±2 kb of each transcriptional start site (TSS) were calculated in each sample based on hg38 reference coordinates. Then, we averaged across multiple TSSs of one gene in each sample. Two-dimensional kernel density was formed based on all genes TSSs H3K4me3 and H3K27me3 signal. This was visualized using the contour plot of the signals. The High/low cutoff for H3K4me3 was set at four standard deviations below the mean value of the H3K4me3-high distribution [42]. Activated, bivalent or repressed status of genes from control to over-expression OC2 samples can be identified in the contours plot. Each red dots represent the gene has OC2 binding peak(s).

### Super-enhancer annotation

Rank Order of Super Enhancers (ROSE) was used to identify enhancers defined as H3K27Ac peaks 2 Kb away from all TSSs. Following stitching enhancer elements clustered within a distance of 12.5 Kb, all super-enhancers in each sample were identified using a cutoff at the inflection point (tangent slope = 1) based on the ranking order of all typical-enhancers and super-enhancers [79]. ROSE also provided closest genes of each typical-enhancer and super-enhancer, which can be used for functional analysis of these super-enhancers.

### Statistics and reproducibility

Statistically significant data for *in vitro* and *in vivo* assays were assessed by unpaired two-tailed Student’s t-test or Wilcoxon two-tailed rank-sum test, unless otherwise noted. Tests for difference in the case of more than two groups were performed using one-way ANOVA where applicable using the Dunnett’s post hoc test, unless otherwise indicated. We used GraphPad Prism and the R (v.3.5, http://www.r-project.org/) for all statistical tests.

