## Supplemental Figure 1 to 7 for "ONECUT2 Activates Diverse Resistance Drivers of Androgen Receptor-Independent Heterogeneity in Prostate Cancer" (from another version of this preprint: version 1, 2023-10-01)

Supplementary Figure 1

(A) OC2 ISH intensity level in each DSP class.
(B) OC2 IHC staining score in specimens with low (<2) and high (⩾2) ISH score.
(C) Representative IHC staining of OC2 in CRPC specimens with distinct features.
(D) Heatmap of AR/NE/Stem signature genes in DSP cohorts. NEPC specimens contain both Stem+ and Stem- phenotypes.
(E) Epithelial cell annotation based on *EPCAM, KRT8* and *CDH1* markers.
(F) Unsupervised clustering using TF activity from the DoRothEA database shows 4 distinct subtypes.
(G) Hallmark enrichment analysis shows distinct pathway activation in each subtype.
(H) In CSPC samples, M1 status displays higher OC2 activity compared to M0 status (scRNA-seq (GSE141445, GSE157703, and GSE137829) and bulk RNA-seq datasets (GSE147493).
(I) UMAP plot of OC2 activity along the pseudotime trajectory.
(J) UMAP plot illustrating PSCA and ASCL1 expression levels in OC2-expressing cells.

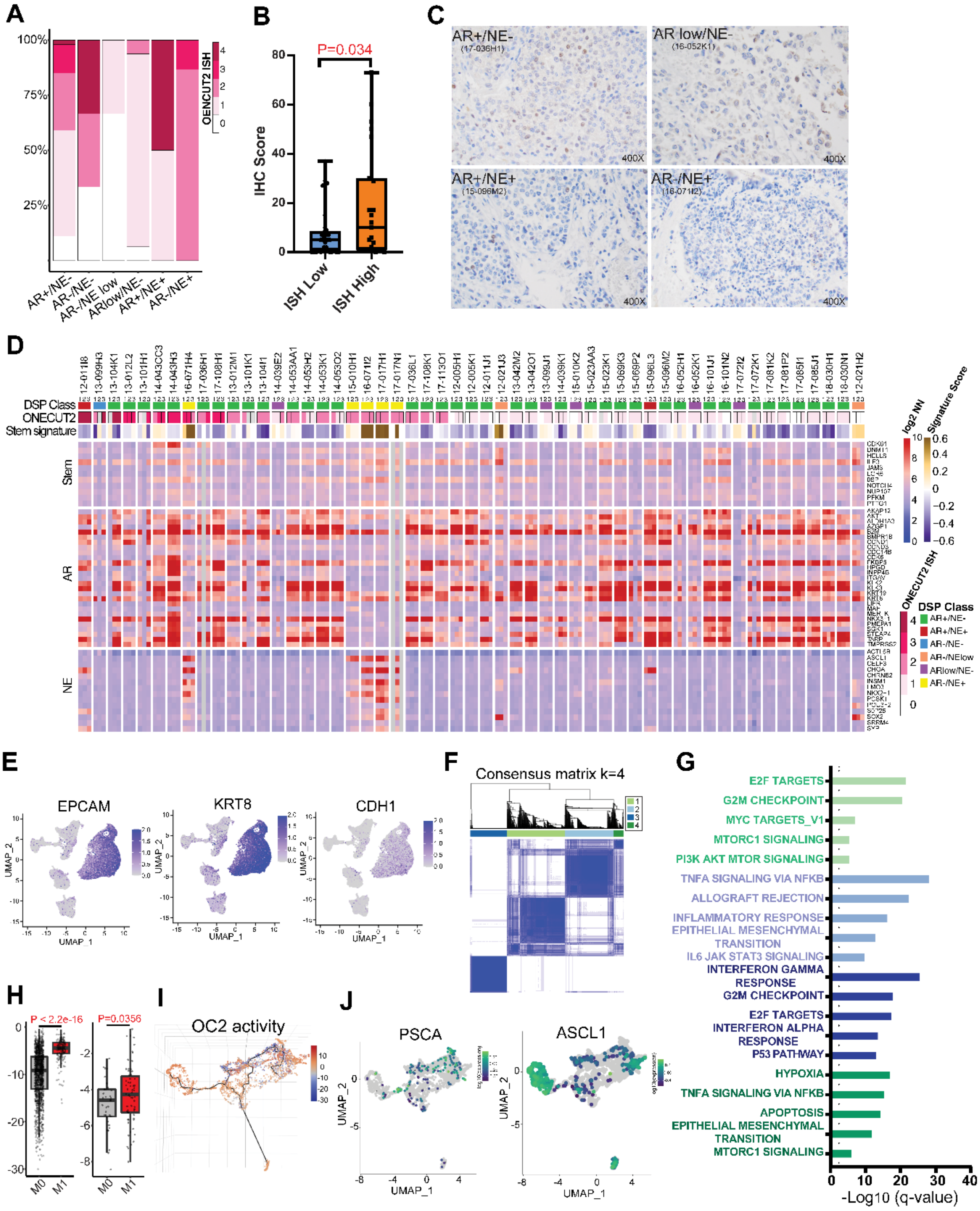
A
100%
75%
50%
25%
AR+/NE-
AR-/NE-
AR-/NE low
ARlow/NE-
AR+/NE+
AR-/NE+
OENCUT2 ISH
B
P=0.034
IHC Score
ISH Low
ISH High
C
AR+/NE-
AR low/NE-
AR+/NE+
AR-/NE+
400X
D
DSP Class
ONECUT2
Stem signature
Stem
AR
NE
log2 NN
Signature Score
ONECUT2 ISH
AR+/NE-
AR+/NE+
AR-/NE-
AR-/NElow
ARlow/NE-
AR-/NE+
E
EPCAM
KRT8
CDH1
UMAP_1
UMAP_2
F
Consensus matrix k=4
G
E2F TARGETS
G2M CHECKPOINT
MYC TARGETS_V1
MTORC1 SIGNALING
PI3K AKT MTOR SIGNALING
TNFA SIGNALING VIA NFKB
ALLOGRAFT REJECTION
INFLAMMATORY RESPONSE
EPITHELIAL MESENCHYMAL TRANSITION
IL6 JAK STAT3 SIGNALING
INTERFERON GAMMA RESPONSE
G2M CHECKPOINT
E2F TARGETS
INTERFERON ALPHA RESPONSE
P53 PATHWAY
HYPOXIA
TNFA SIGNALING VIA NFKB
APOPTOSIS
EPITHELIAL MESENCHYMAL TRANSITION
MTORC1 SIGNALING
-Log10 (q-value)
H
P < 2.2e-16
P=0.0356
M0
M1
I
OC2 activity
J
PSCA
ASCL1
UMAP_1
UMAP_2

Supplementary Figure 2.

(A) Highest abundant proteins in the OC2 interactome list (Average Counts in OC2 OE, Vec Con serve as negative control group).
(B) New enhancer looped to OC2 promoter regions under OC2 OE conditions.
(C) ATAC-seq signaling profiling of LNCaP Vec Con and OC2 OE cells across the genome.
(D) Distribution of accessible regions within promoters and inter-genic regions in LNCaP Vec Con and OC2 OE cells.
(E) Mass-spectrometry peptide counts for CRC families.

**A** Top 20 abundant proteins

| Protein names | Gene names | Average Counts |
|---|---|---|
| Neuroblast differentiation-associated protein AHNAK | AHNAK | 105 |
| DNA-dependent protein kinase catalytic subunit | PRKDC | 97.5 |
| Pre-mRNA-processing-splicing factor 8 | PRPF8 | 62.5 |
| E3 SUMO-protein ligase RanBP2 | RANBP2 | 62.5 |
| ATP-dependent RNA helicase A | DHX9 | 55.5 |
| Structural maintenance of chromosomes protein 3 | SMC3 | 54 |
| Structural maintenance of chromosomes protein 1A | SMC1A | 49.5 |
| Nuclear mitotic apparatus protein 1 | NUMA1 | 49 |
| SWI/SNF-related matrix-associated actin-dependent regulator of chromatin subfamily A member 5 | SMARCA5 | 47 |
| Spectrin alpha chain, non-erythrocytic 1 | SPTAN1 | 45 |
| Spectrin beta chain, non-erythrocytic 2 | SPTBN2 | 44 |
| Double-stranded RNA-specific adenosine deaminase | ADAR | 40.5 |
| Nuclear pore complex protein Nup153 | NUP153 | 40.5 |
| Myosin-9 | MYH9 | 40 |
| U5 small nuclear ribonucleoprotein 200 kDa helicase | SNRNP200 | 39.5 |
| Splicing factor 3B subunit 1 | SF3B1 | 38.5 |
| Probable ATP-dependent RNA helicase DDX5 | DDX5 | 38.5 |
| Antigen KI-67 | MKI67 | 38 |
| Poly [ADP-ribose] polymerase 1 | PARP1 | 37.5 |
| Homeobox protein cut-like 1 | CUX1 | 37.5 |

**C**

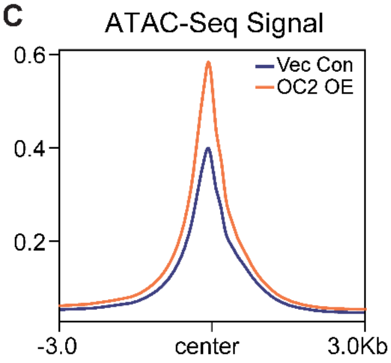


**D**

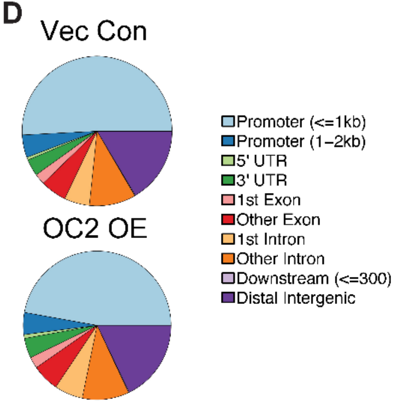


**B**

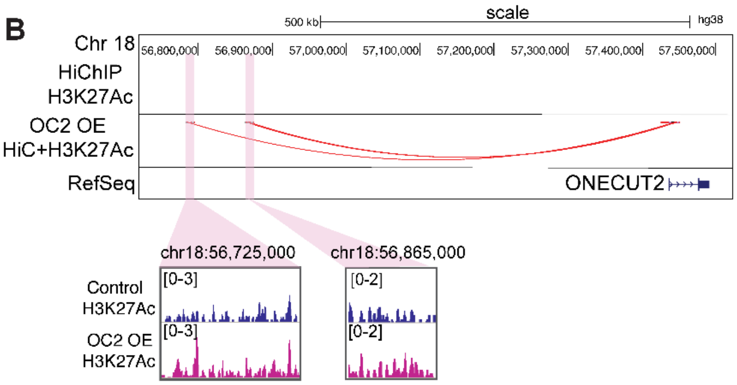


**E**

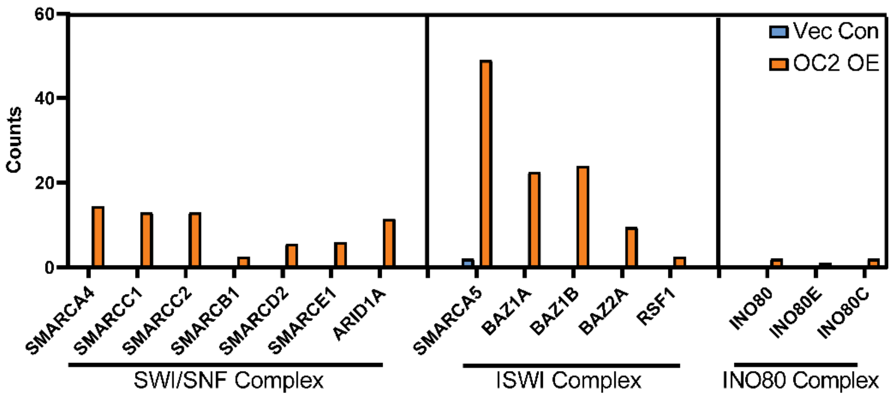

Supplementary Figure 3.

(A) RNA-seq score of the top candidate TFs driven by OC2.

(B) ATAC-seq score (Motif displacement score) of the top candidate TFs driven by OC2.

(C) Integrated Z-score with the ranking of candidate TFs. Positive z-score represents OC2 activated TFs, Negative z-score represents OC2-suppressed TFs. (**Refer to methods**)

(D) Fold enrichment score with the ranking of candidate TFs. (**Refer to methods**)

(E) The cutoff of H3K4me3 signal in LNCaP Vec Con and OC2 OE cells fitted in a two normal distribution model.

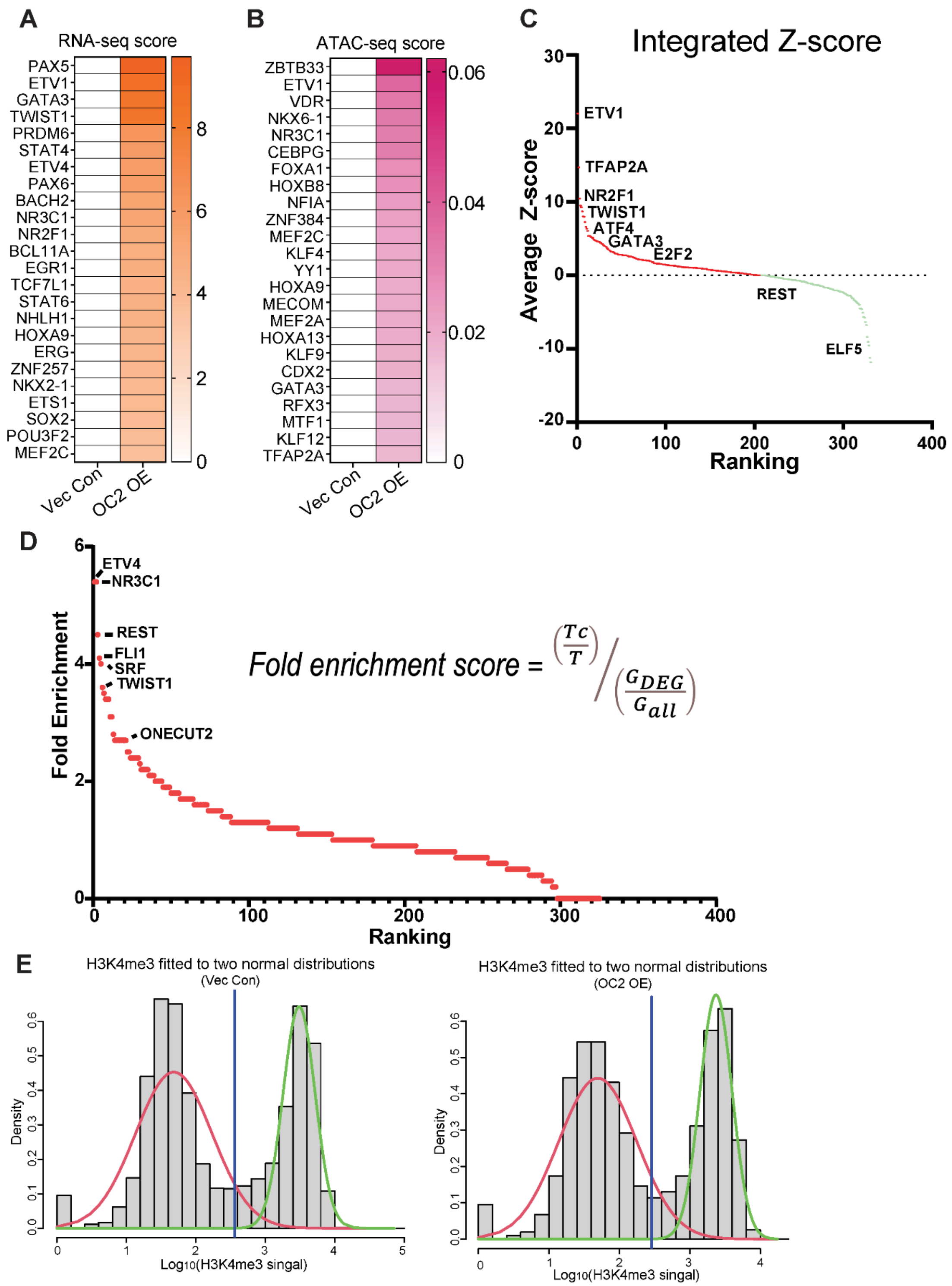

A
RNA-seq score
PAX5
ETV1
GATA3
TWIST1
PRDM6
STAT4
ETV4
PAX6
BACH2
NR3C1
NR2F1
BCL11A
EGR1
TCF7L1
STAT6
NHLH1
HOXA9
ERG
ZNF257
NKX2-1
ETS1
SOX2
POU3F2
MEF2C
Vec Con
OC2 OE
B
ATAC-seq score
ZBTB33
ETV1
VDR
NKX6-1
NR3C1
CEBPG
FOXA1
HOXB8
NFIA
ZNF384
MEF2C
KLF4
YY1
HOXA9
MECOM
MEF2A
HOXA13
KLF9
CDX2
GATA3
RFX3
MTF1
KLF12
TFAP2A
C
Integrated Z-score
Average Z-score
Ranking
ETV1
TFAP2A
NR2F1
TWIST1
ATF4
GATA3
E2F2
REST
ELF5
D
Fold Enrichment
Ranking
ETV4
NR3C1
REST
FLI1
SRF
TWIST1
ONECUT2
Fold enrichment score = $\left(\frac{Tc}{T}\right) / \left(\frac{G_{DEG}}{G_{all}}\right)$
E
H3K4me3 fitted to two normal distributions
(Vec Con)
Density
Log10(H3K4me3 singal)
H3K4me3 fitted to two normal distributions
(OC2 OE)
Density
Log10(H3K4me3 singal)

Supplementary Figure 4.
(A) OC2 suppressed AR downstream targets in LNCaP cells.
(B) AR response hallmark genes were suppressed with OC2 overexpression.
(C) Knockdown of OC2 in LNCaP OC2 OE cells suppressed AR target genes.
(D) Suppression of OC2 by removal of cumate suppressed GR expression in LAPC4 cells.
(E) OC2 bound to the *NR3C1* promoter region.

Supplementary Figure 4.

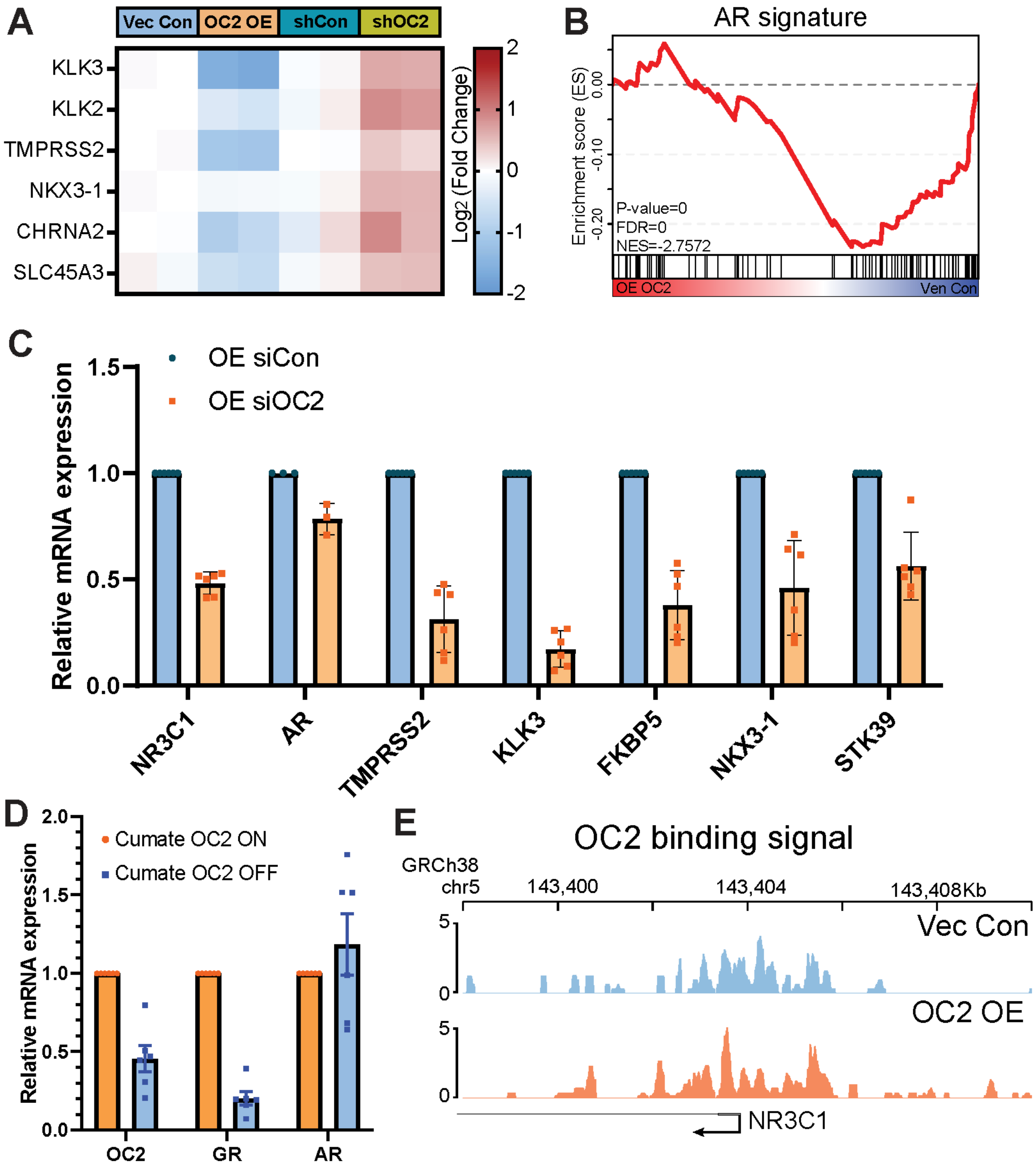

Supplementary Figure 5.
(A) Upregulation of NE markers in LAPC4 cells with OC2 overexpression.
(B) Neural stem lineage driver ASCL1 is activated by OC2.
(C) OC2 signature genes derived in LNCaP cells identify PDXs with NE features comparable to the Beltran NEPC signature [31].
(D) Super-enhancer calling using the ROSE algorithm in both Vec Con and OC2 OE cells (N=2).
(E) Gene set enrichment of SE-associated genes in LNCaP Vec Con and OC2 OE cells.
(F) Representative specific super-enhancer regions are shown.
(G) Heatmap of SE-associated genes validated in SU2C cohort.
(H) OC2 upregulates SRRM4 in 22Rv1 cells.
(I) Suppression of OC2 did not reverse the NE phenotype in OC2 OE cells.

Supplementary Figure 5.

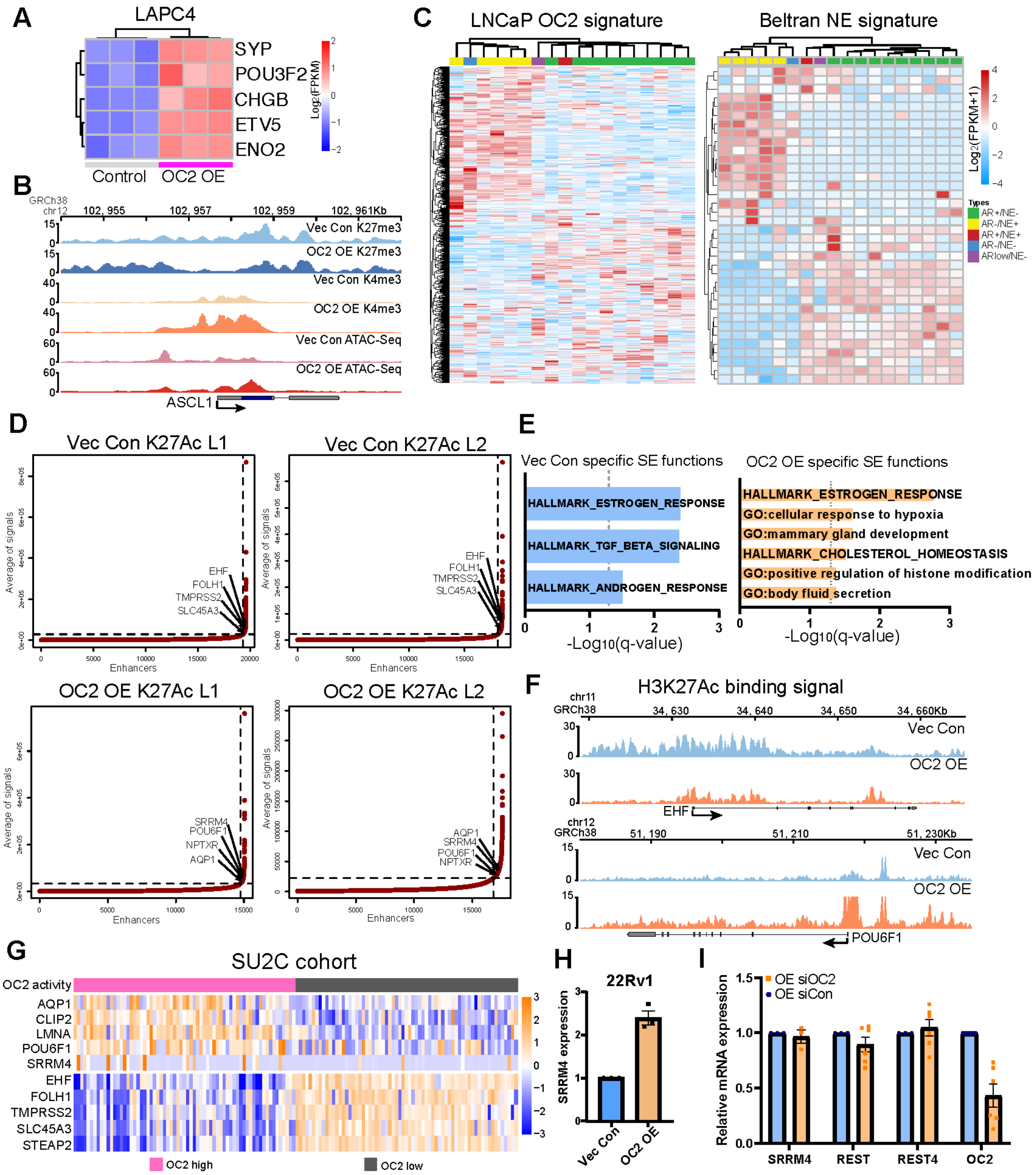

Supplementary Figure 6.

(A) OC2, AR and FOXA1 bind directly to the *UGT2B15/17* promoter region as seen by CUT&RUN sequencing.

(B) Predicted OC2 binding region with the scanmotifgenomewide function using the Hypergeometric Optimization of Motif EnRichment (HOMER) package [39].

(C) Enzalutamide only activates UGT2B15 in LNCaP Vec Con cells.

(D) OC2 binding sites are conserved across multiple UGT2B family members.

Supplementary Figure 6.

**A**

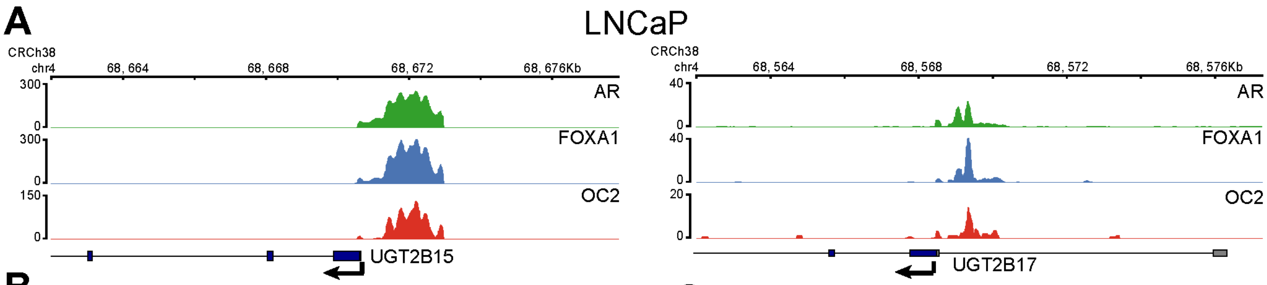


**B**

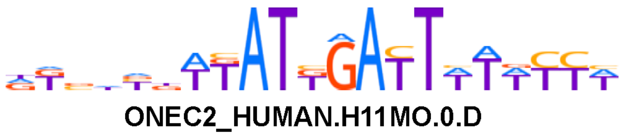


| Chr | Start | End | Log Odds Score | Strand |
|---|---|---|---|---|
| Chr4 | 69536404 | 69536424 | 12 | - |
| Chr4 | 69535296 | 69535316 | 11 | + |
| Chr4 | 69542158 | 69542178 | 9 | - |
| Chr4 | 69543186 | 69543206 | 9 | + |
| Chr4 | 69543192 | 69543212 | 9 | + |

**C**

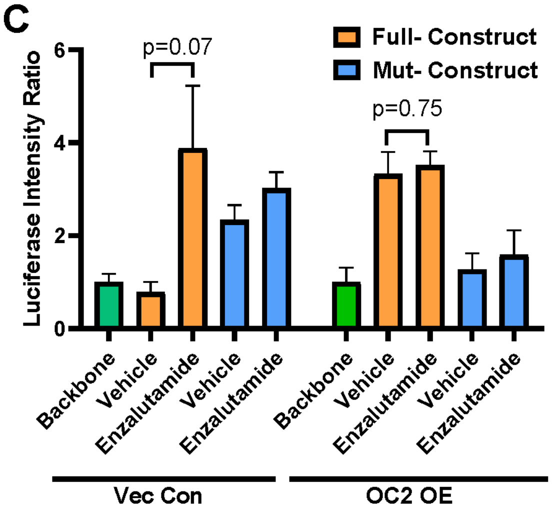


**D**

OC2 binding site

| | | OC2 binding site | |
|---|---|---|---|
| **UGT2B15** | GCAGTTATATTTTAA | CTTGATTGATTTTT | CCTCAGATATAAGTATGAGAAATGACAGAAAGAAACAAC |
| UGT2B17 | GCAGTTATATTTTAA | CTTGATTGATTTTT | CCTCAGATATAAGTATGAGAAATGACAGAAAGAAACAAC |
| UGT2B10 | G-AGTTATGTTTTAA | CTTGATTGATTTAT | CTCTGTATATAAGTATGAGAAAGAAACAGTGACTGGAAA |
| UGT2B11 | G-AGTTATATTTTAA | CTTGATTGATTTAT | CTCTGTATATAAGTATGAGAAAGAAACAGTGACTGGAAA |
| UGT2B28 | TAATTCCTCAAAATA | CTTGATTTTCTCTC | TTTAATATTTGGTACCAATTCTTTAGTAGTGCCTTAGTG |

Supplementary Figure 7.
(A)  Suppression of REST and upregulation of OC2 by enzalutamide in LNCaP cells.
(B) Heatmap of gene expression in control, enzalutamide treatment alone and combined treatment with OC2 inhibitor.
(C) Gene set enrichment of suppressed genes in combined treatment vs. enzalutamide alone.
(D) OC2 activated Stem and NE signature genes. The suppressed binding intensity of AR in OC2 OE cells on lineage driver SOX2 and BRN2 is shown.
(E). Heatmap of AR, Stem, and NE signature gene expression in control, enzalutamide treatment alone and combined treatment with OC2 inhibitor.
(F). Workflow for identifying FOXA1-mediated chromatin remodeling involved in the combined treatment condition.
(G) Activation of AR signaling with OC2 inhibitor used alone seen with GFP tagged-AR response system.
(H) Pre-treatment of LNCaP cells with OC2 inhibitor 3 days prior to enzalutamide induced extensive cell death.
(I) Suppressed hallmarks in pre-treatment combination treatment (panel G) compared with enzalutamide alone.
Supplementary Figure 7.

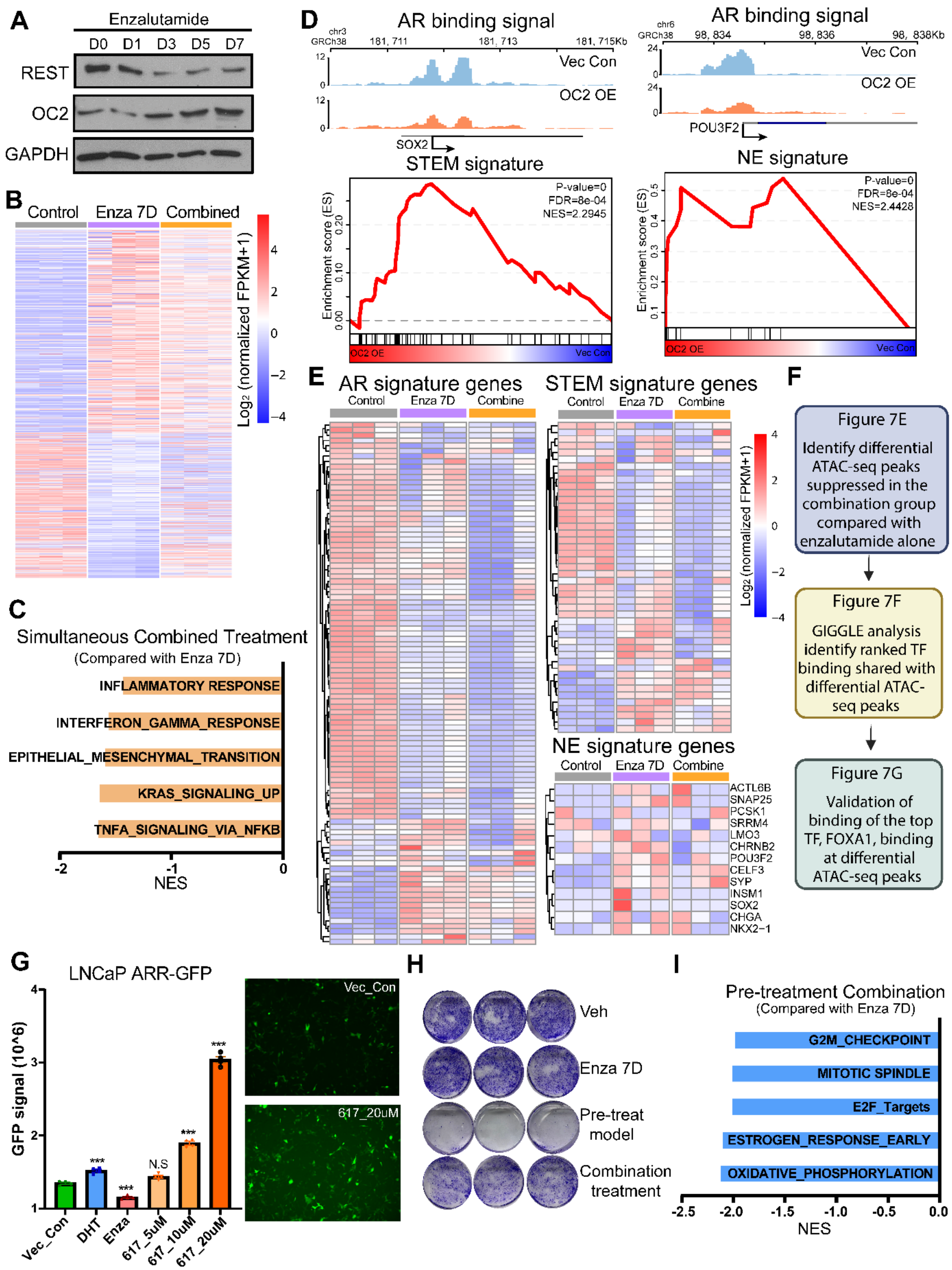
A
Enzalutamide
D0 D1 D3 D5 D7
REST
OC2
GAPDH
B
Control Enza 7D Combined
Log2 (normalized FPKM+1)
C
Simultaneous Combined Treatment
(Compared with Enza 7D)
INFLAMMATORY RESPONSE
INTERFERON_GAMMA_RESPONSE
EPITHELIAL_MESENCHYMAL_TRANSITION
KRAS_SIGNALING_UP
TNFA_SIGNALING_VIA_NFKB
NES
D
AR binding signal
Vec Con
OC2 OE
SOX2
POU3F2
STEM signature
P-value=0
FDR=8e-04
NES=2.2945
NE signature
P-value=0
FDR=8e-04
NES=2.4428
Enrichment score (ES)
E
AR signature genes
STEM signature genes
NE signature genes
Control Enza 7D Combine
ACTL6B
SNAP25
PCSK1
SRRM4
LMO3
CHRNB2
POU3F2
CELF3
SYP
INSM1
SOX2
CHGA
NKX2−1
F
Figure 7E
Identify differential ATAC-seq peaks suppressed in the combination group compared with enzalutamide alone
Figure 7F
GIGGLE analysis identify ranked TF binding shared with differential ATAC-seq peaks
Figure 7G
Validation of binding of the top TF, FOXA1, binding at differential ATAC-seq peaks
G
LNCaP ARR-GFP
GFP signal (10^6)
Vec_Con DHT Enza 617_5uM 617_10uM 617_20uM
N.S
Vec_Con
617_20uM
H
Veh
Enza 7D
Pre-treat model
Combination treatment
I
Pre-treatment Combination
(Compared with Enza 7D)
G2M_CHECKPOINT
MITOTIC SPINDLE
E2F_Targets
ESTROGEN_RESPONSE_EARLY
OXIDATIVE_PHOSPHORYLATION
NES
